# Schizophrenia-associated DNA methylation differences in the cortex are neuron-specific

**DOI:** 10.64898/2026.07.08.737176

**Authors:** Eilis Hannon, Emma M Walker, Barry Chioza, Joe Burrage, Georgina E T Blake, Madaleine Sharp, Ann Babtie, Marty Frith, Nicholas Clifton, Leonard Schalkwyk, Emma L Dempster, Jonathan Mill

## Abstract

Schizophrenia is a complex neuropsychiatric disorder in which genetic risk is thought to converge on cell type-specific regulatory mechanisms in the brain. We performed a cell type-resolved epigenome-wide association study (EWAS) of schizophrenia using fluorescence-activated nuclei sorting (FANS) to isolate neuron-enriched (NeuN+), oligodendrocyte-enriched (SOX10+) and other glial-enriched (NeuN−/SOX10−) nuclei populations alongside total prefrontal cortex nuclei fractions from 216 donors (104 schizophrenia cases and 112 controls). We identified 16 differentially methylated positions (DMPs) in neuron-enriched nuclei at experiment-wide significance and more than 400 additional neuronal DMPs at a discovery threshold. In contrast, no significant associations were identified in oligodendrocyte-enriched, glial-enriched or total nuclei fractions, demonstrating that schizophrenia-associated cortical methylomic variation is highly neuron-specific and largely masked in bulk tissue analyses. Neuronal DMPs exhibited a significant bias towards hypomethylation in schizophrenia and were enriched at loci implicated by genetic studies, including *CACNA1C, CACNA1G* and *TRIO*. Pathway analyses implicated genes involved in neurodevelopment, cell adhesion, synapse organisation, neurotransmission and synaptic plasticity. Schizophrenia-associated DNA methylation signatures identified in prefrontal cortex neurons showed correlated effects in neuronal nuclei isolated from the hippocampus and striatum, indicating partial conservation of disease-associated epigenetic alterations across brain regions. Together, these findings provide strong evidence for widespread neuron-specific epigenetic dysregulation in schizophrenia and highlight the importance of cell type-resolved approaches for elucidating the molecular mechanisms underlying psychiatric disease.

## INTRODUCTION

Schizophrenia is a severe psychiatric disorder marked by recurrent psychosis, negative symptoms and cognitive impairment^1^. Risk for developing schizophrenia is highly polygenic, involving hundreds of common variants alongside rare copy-number and coding changes^2–4^. Evidence suggests that genetic risk for schizophrenia operates through the disruption of neuron-specific gene networks. Single-cell transcriptomic studies demonstrate enrichment of schizophrenia risk genes in cortical excitatory and inhibitory neurons^5^, while genetic risk variants are preferentially localised to neuron-specific regulatory elements, consistent with non-coding mechanisms acting in neuronal populations^6,7^. In addition to DNA sequence variation, environmental factors including prenatal adversity, infection, and cannabis exposure are also known to shape risk trajectories. Epigenetic mechanisms, particularly DNA methylation, provide a molecular interface through which genetic and environmental influences modulate gene regulation in specific cell types across development and into adulthood^8^.

Previous epigenome-wide association studies (EWAS) of schizophrenia have focused on profiling DNA methylation in either peripheral blood^9^ or bulk post-mortem cortex tissue^10,11^. Although these studies have identified schizophrenia-associated DNA methylation differences, interpretation has been limited by the cellular complexity of the human cortex. Variation in cellular composition can confound bulk tissue analyses, while disease-associated changes occurring within specific cell populations are likely to be diluted or obscured. Statistical deconvolution approaches can partially control for cellular heterogeneity^12^, but cannot directly characterise cell type-specific DNA methylation differences. Emerging studies separating neuronal and non-neuronal nuclei suggest that schizophrenia-associated epigenetic variation is enriched in neurons^13^, but these analyses are limited by modest sample sizes and have only compared neuronal and non-neuronal nuclei.

In this study, we performed a large cell-type-resolved study of DNA methylation differences associated with schizophrenia in human PFC. Using a fluorescence-activated nuclei sorting (FANS) approach developed by our group^14^, we isolated neuron-enriched (NeuN+), oligodendrocyte-enriched (SOX10+) and other glial-enriched (NeuN−/SOX10−) nuclei populations, alongside matched total nuclei fractions, from 216 donors (104 schizophrenia cases and 112 non-psychiatric controls). We then performed cell type-specific EWAS analysis to identify schizophrenia-associated DNA methylation differences across cortical cell populations. By characterising the cellular specificity of schizophrenia-associated methylomic variation, this study provides new insights into the neuronal regulatory mechanisms linking genetic risk to cortical circuit dysfunction in schizophrenia.

## RESULTS

### Overview of dataset

We used a FANS method developed by our group to isolate highly-enriched populations of neuron-enriched (NeuN+), oligodendrocyte-enriched (SOX10+) and other glial cell (NeuN-/SOX10-) nuclei (**Supplementary Figure 1**) from PFC tissue dissected from 216 donors (n = 104 schizophrenia cases, n = 112 non-psychiatric controls) (**Supplementary Table 1**). After FANS, DNA methylation was quantified across the genome for each nuclei fraction alongside a ‘total’ nuclei fraction reflecting bulk PFC tissue from each donor using the Illumina EPIC HumanMethylation microarray (see **Methods**). Following stringent pre-processing, our final dataset included DNA methylation data for 779,868 sites (762,508 autosomal sites and 17,360 on the X chromosome) across 653 nuclei fractions (NeuN+: cases = 77, controls = 84; SOX10+: cases = 69, controls = 84; NeuN-/SOX10-: cases = 65, controls = 77; total nuclei: cases = 96, controls = 101). The efficiency and specificity of nuclei purification were confirmed experimentally using single-nucleus RNA sequencing performed on sorted fractions from four donors demonstrating the expected enrichment of cell type-specific transcriptional signatures within each population (**Supplementary Figure 2)**. FANS purification was further validated using a cellular deconvolution approach for different brain cell types developed by our group^12^ which confirmed near-perfect enrichment of each nuclei population (**Supplementary Figure 3)** with no difference in purity between cases and controls for each fraction (NeuN+: cases = 0.970, controls = 0.975, P = 0.13; SOX10+: cases = 1.00, controls = 0.996, P = 0.66; NeuN-/SOX10-: cases = 0.946, controls = 0.934, P = 0.62). Unsupervised analyses, including principal component analysis (PCA) and hierarchical clustering, revealed marked separation of nuclei fractions according to cell type (**Supplementary Figure 4**), confirming the strong cell type specificity of cortical DNA methylation patterns and emphasising the importance of cell type-resolved profiling approaches for identifying schizophrenia-associated epigenetic variation.

### Cell type-resolved EWAS identifies widespread neuronal hypomethylation in schizophrenia

To identify schizophrenia-associated differences in DNA methylation, we performed epigenome-wide association studies (EWAS) separately within each nuclei fraction (NeuN+, SOX10+, NeuN-/SOX10- and ‘total’), fitting linear regression models controlling for age, sex and brain bank (see **Methods**). Quantile-quantile (QQ) plots show little evidence for test statistic inflation across all nuclei populations (**Supplementary Figure 5**), supporting the robustness of the analytical approach. Across all analyses, we identified 16 differentially methylated positions (DMPs) associated with schizophrenia at a stringent experiment-wide significance threshold (P < 9.00E-8) (**Figure 1A** and **Table 1**). All 16 DMPs were identified exclusively in neuronal nuclei, with no experiment-wide significant loci detected in oligodendrocyte, other glial or total nuclei fractions. This strong neuronal enrichment of schizophrenia DMPs was also evident at a more relaxed discovery threshold (P < 1.00E-4), where 428 of 505 DMPs (84.8%) were identified in NeuN+ nuclei, compared with only 23 in SOX10+ nuclei, 14 in NeuN−/SOX10− nuclei and 40 in total nuclei fractions (**Supplementary Tables 2-5**). Together, these findings indicate that schizophrenia-associated methylomic variation in the cortex is highly cell type-specific and is largely masked in analyses of bulk tissue. Comparison of absolute mean effect sizes across different nuclei fractions further demonstrated the neuronal specificity of schizophrenia-associated differences (**Figure 1B**) with the mean effect size at the 428 DMPs identified in the neuron-enriched population being significantly larger than in the other nuclei populations (Wilcoxon signed rank test (paired): NeuN+ vs SOX10+ P = 3.751E-72; NeuN+ vs NeuN-/SOX10-P = 1.059E-71; NeuN+ vs total P = 3.804E-72). Although effect estimates for neuronal DMPs were directionally correlated with those observed in the total nuclei fraction (corr = 0.54), effect sizes were consistently attenuated in bulk tissue, highlighting the increased sensitivity and power afforded by cell type-resolved profiling approaches (**Supplementary Figure 6**). Finally, we formally tested the cell type-specificity of NeuN+ DMPs using a mixed effects model (see **Methods**). Effects at all 16 experiment-wide significant DMPs were cell-type-specific (ANOVA P < 0.05) with the majority (74.3%) of discovery-threshold DMPs also showing significant cell-type-specific effects. These results confirm that there is heterogeneity in effects across cell types and that the identified differences are largely specific to NeuN+ nuclei.

**Figure 1:**
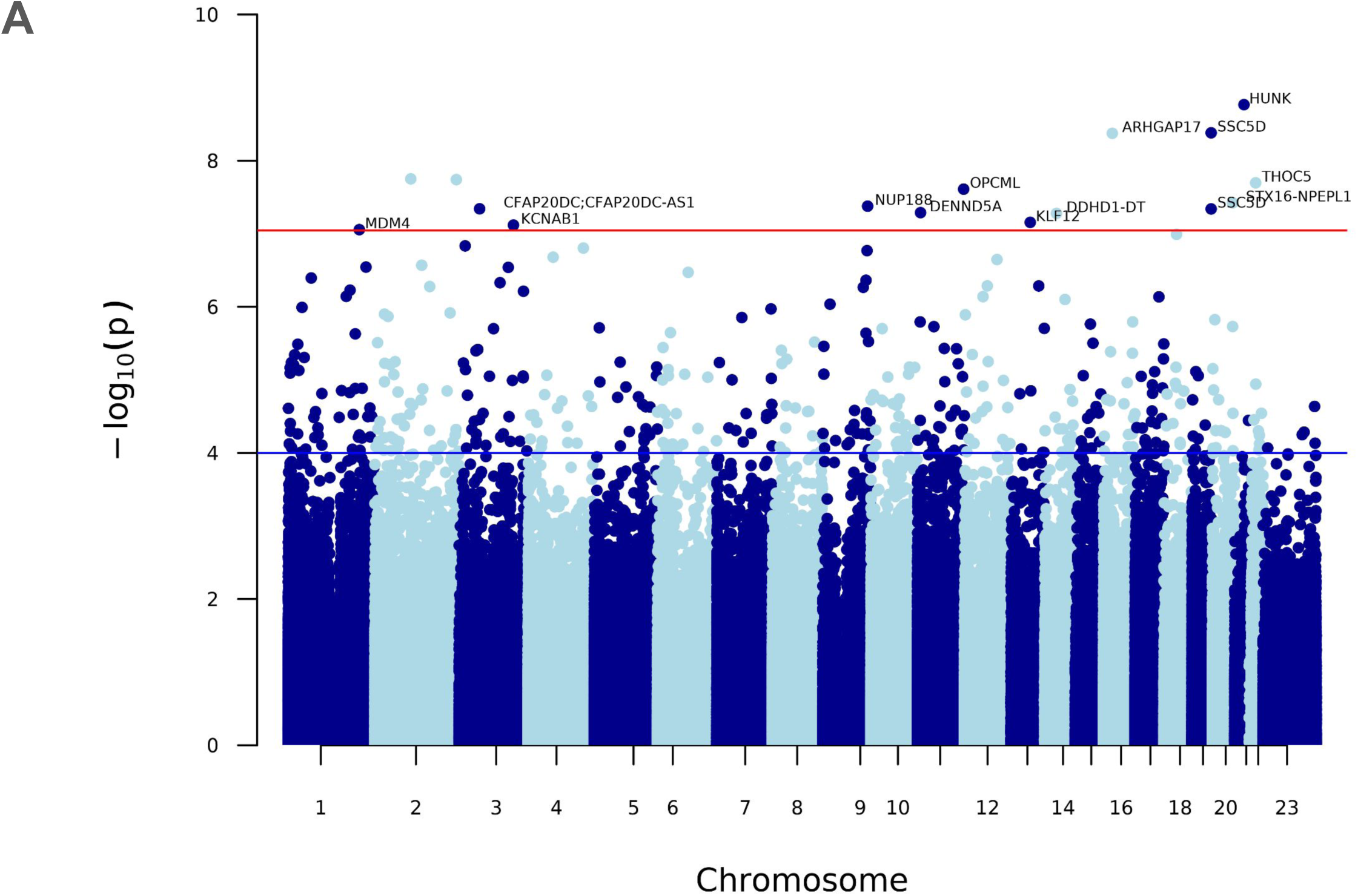
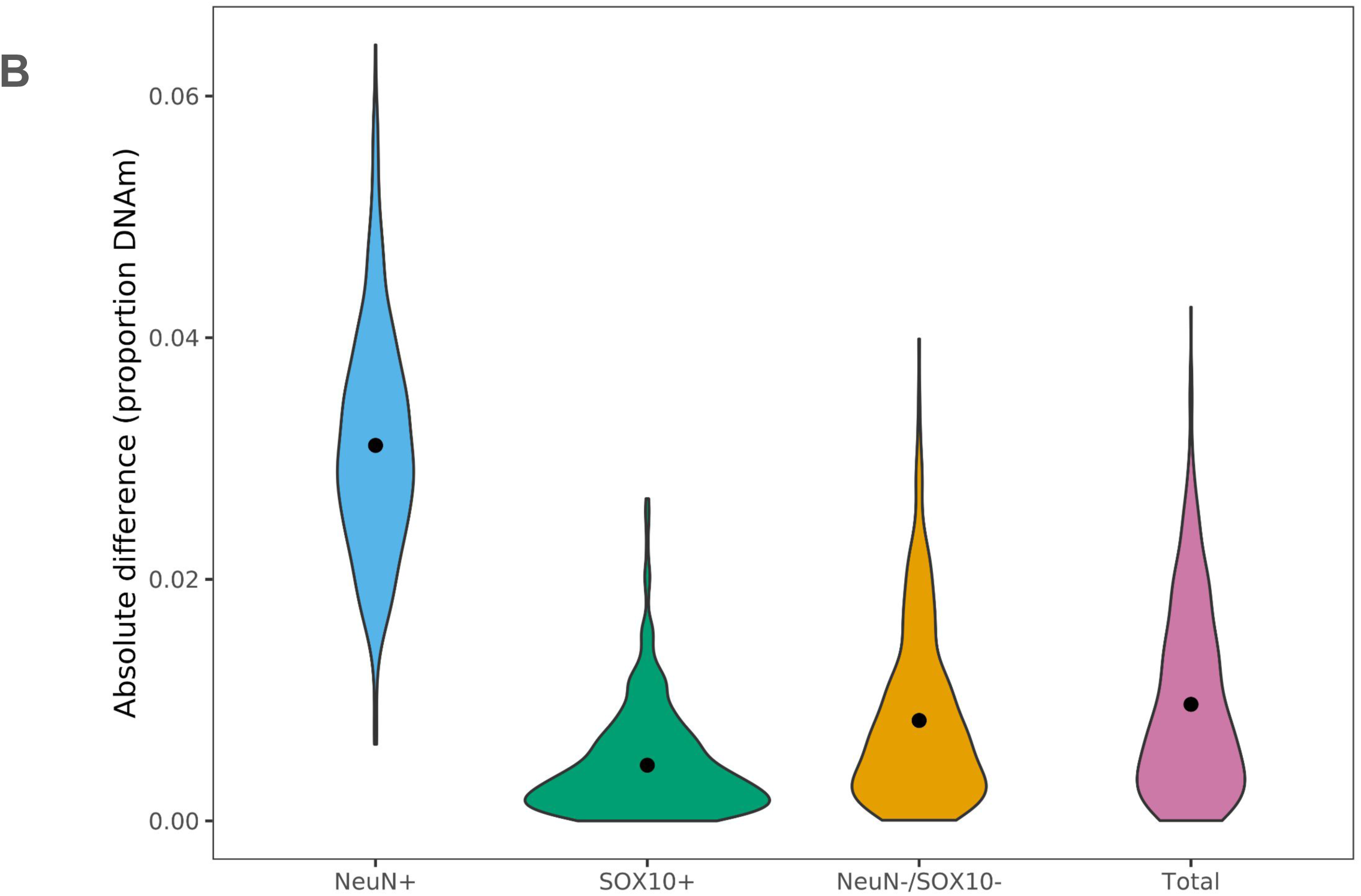
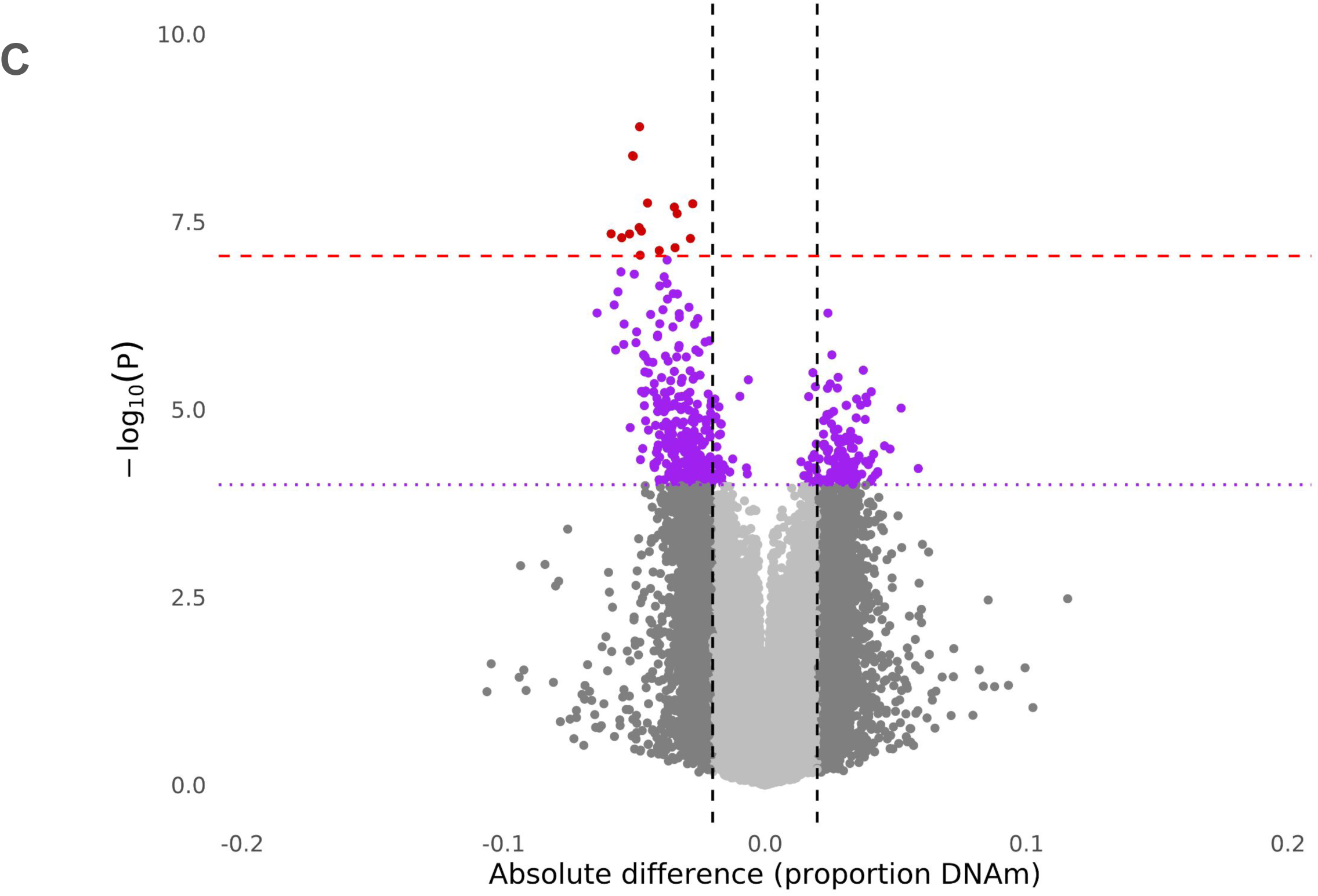
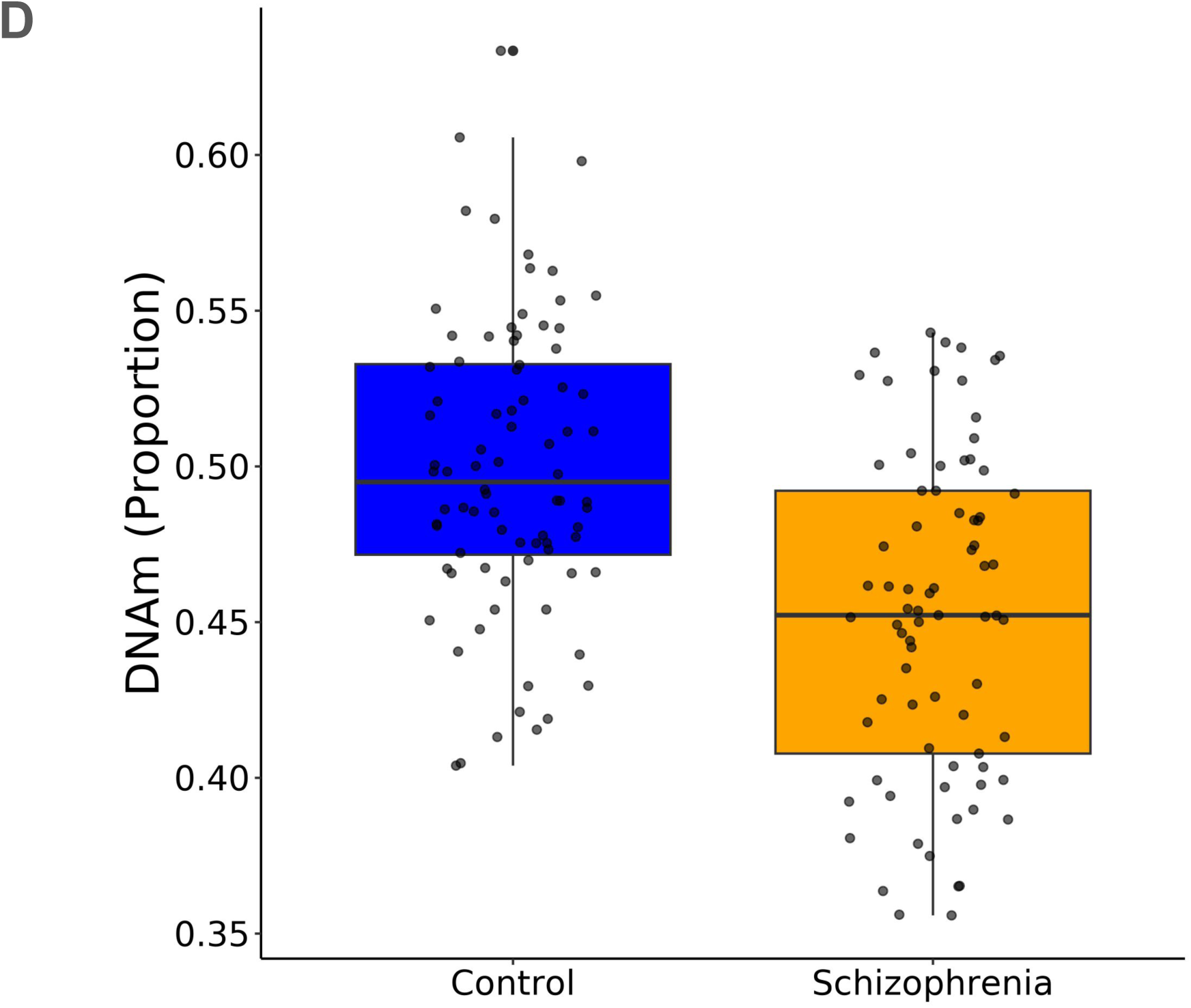
Schizophrenia-associated DNA methylation differences are highly neuron-specific and characterised by widespread hypomethylation. A) Manhattan plot showing the genomic distribution of schizophrenia-associated differentially methylated positions (DMPs) identified in neuron-enriched (NeuN+) nuclei isolated from prefrontal cortex (PFC). B) Comparison of effect sizes for 428 schizophrenia-associated NeuN+ DMPs identified at the discovery threshold (P < 1.00E-4) demonstrates marked neuronal specificity, with significantly larger effects observed in NeuN+ nuclei than in oligodendrocyte-enriched (SOX10+), other glial-enriched (NeuN−/SOX10−) or total nuclei fractions (t test (paired): NeuN+ vs SOX10+ P = 2.871E-198; NeuN+ vs NeuN-/SOX10-P = 4.90E-157; NeuN+ vs total P = 2.14E-187)). C) Schizophrenia-associated neuronal DMPs exhibit a strong directional bias towards hypomethylation. All experiment-wide significant DMPs (red) were hypomethylated in schizophrenia cases, and hypomethylated sites were significantly enriched among discovery-threshold DMPs (purple). D) The strongest schizophrenia-associated DMP was cg08080338, annotated to the gene encoding hormonally up-regulated Neu-associated kinase (*HUNK*). DNA methylation at this site was significantly reduced in schizophrenia cases specifically within NeuN+ nuclei (effect size = 0.048, P = 1.711E-9).

**Table 1:** Differentially methylated positions (DMPs) associated with schizophrenia at experiment-wide significance. Shown are sites at which DNA methylation is associated with schizophrenia at an experiment-wide significance threshold of (P < 9.00E-8). All significant DMPs were identified exclusively within the neuron-enriched (NeuN+) nuclei fraction. Reported statistics include genomic coordinates, annotated gene, effect size and association P-value. Full lists of DMPs identified at the discovery significance threshold (P < 1.00E-4) for each nuclei fraction are provided in **Supplementary Tables 2-5**.

| Site | Case-control difference | P | Chr | Position | Gene |
| --- | --- | --- | --- | --- | --- |
| cg08080338 | -0.04796 | 1.711E-09 | 21 | 31921302 | *HUNK* |
| cg12421899 | -0.05063 | 4.157E-09 | 19 | 55520020 | *SSC5D* |
| cg05583636 | -0.05038 | 4.223E-09 | 16 | 24994441 | *ARHGAP17* |
| cg26664243 | -0.04492 | 1.774E-08 | 2 | 103435095 |  |
| cg14160339 | -0.02762 | 1.816E-08 | 2 | 234289936 |  |
| cg00848123 | -0.03468 | 2.010E-08 | 22 | 29521240 | *THOC5* |
| cg18944144 | -0.0336 | 2.454E-08 | 11 | 132803604 | *OPCML* |
| cg17952138 | -0.04812 | 3.768E-08 | 20 | 58682920 | *STX16-NPEPL1* |
| cg16153363 | -0.04726 | 4.195E-08 | 9 | 128973087 | *NUP88* |
| cg04246167 | -0.05886 | 4.557E-08 | 3 | 58998739 | *CFAP20DC* |
| cg25778661 | -0.0518 | 4.576E-08 | 19 | 55520006 | *SSC5D* |
| cg13173567 | -0.0548 | 5.144E-08 | 11 | 9150743 | *DENND5A* |
| cg01070948 | -0.0285 | 5.269E-08 | 14 | 53614109 | *DDHD1-DT* |
| cg03356288 | -0.03438 | 6.984E-08 | 13 | 74064946 | *KLF12* |
| cg19555318 | -0.04045 | 7.616E-08 | 3 | 156181835 | *KCNAB1* |
| cg14724853 | -0.04775 | 8.790E-08 | 1 | 204521013 | *MDM4* |

In addition to their strong neuronal specificity, schizophrenia-associated DMPs showed a striking directional bias towards hypomethylation in schizophrenia cases. All 16 experiment-wide significant neuronal DMPs were hypomethylated in schizophrenia, and hypomethylated sites were significantly enriched amongst discovery-threshold neuronal DMPs (291 hypomethylated versus 137 hypermethylated sites, binomial P = 3.81E-14, Figure 1C). Together, these findings indicate that schizophrenia-associated cortical methylomic variation is dominated by widespread neuronal DNA hypomethylation.

Because the NeuN+ fraction comprises a mixture of neuronal subtypes, we assessed whether variation in neuronal subtype composition could influence the observed schizophrenia-associated DNA methylation differences. Using a publicly available DNA methylation reference dataset generated from SOX6+ inhibitory (GABAergic) neuronal nuclei15, we estimated the proportion of SOX6+ neurons within each NeuN+ sample using CETYGO16. We then repeated the neuronal EWAS including estimated SOX6+ neuronal proportion as an additional covariate. Adjustment for SOX6+ neuronal abundance had negligible impact on the results, with effect size estimates from models with and without this covariate showing near-perfect concordance (r = 0.996, Supplementary Figure 7). These findings indicate that the schizophrenia-associated DNA methylation differences identified in NeuN+ nuclei are unlikely to be driven by variation in the relative abundance of inhibitory neurons across samples. Association statistics from the adjusted model are provided in Supplementary Table 2.

The most significant neuronal DMP was cg08080338 (effect size = 0.048, P = 1.711E-9), located in intron 1 of the gene encoding hormonally up-regulated Neu-associated kinase (*HUNK*) (**Figure 1D** and **Supplementary Figure 8**), a protein kinase involved in cell polarity, migration and survival with a known role in neurodevelopment via regulation of actin cytoskeleton dynamics and trafficking^17^. Schizophrenia cases exhibited significant hypomethylation at this site specifically within neuronal nuclei, with no significant differences detected in the other cell-type-enriched nuclei populations (**Supplementary Figure 9**). We independently validated this association using targeted bisulfite pyrosequencing in purified NeuN+ nuclei samples from a subset of donors (n = 32), confirming significant schizophrenia-associated hypomethylation across the surrounding amplicon region (amplicon average P = 0.0017) in addition to the site corresponding to cg08080338 (CpG2 in the pyrosequencing assay, P = 0.0194) (**Supplementary Figure 10**). Other significant DMPs mapped to genes involved in neuronal signalling, synaptic organisation and intracellular trafficking, including *ARHGAP17* (effect size = 0.050, P = 4.22E-9), *OPCML* (effect size = 0.034, P = 2.45E-8), *DENND5A* (effect size = 0.055%, P = 5.14E-8) and *KCNAB1* (effect size = 0.040, P = 7.62E-8). Notably, two significant DMPs were identified within 14 bp of each other between the *SSC5D* and *SBK2* genes on chromosome 19 (cg12421899 and cg25778661, **Supplementary Figure 11**), suggesting a localised differentially methylated region.

### Neuronal schizophrenia-associated DMPs converge with genetic risk loci and synaptic pathways

Large-scale genome-wide association and exome sequencing studies have robustly implicated both common and rare genetic variation in schizophrenia susceptibility. We therefore tested whether schizophrenia-associated neuronal DMPs were enriched at genes implicated by (i) rare coding variant analyses from the SCHEMA consortium (n = 31 genes)^3^ and (ii) high-confidence common variant loci identified by the Psychiatric Genomics Consortium (PGC) (n = 108 genes)^4^ (see Methods). Among discovery-threshold neuronal DMPs, nine sites mapped to five PGC-implicated genes (*OPCML* (cg18944144, P = 2.45E-8; cg09531376, P = 9.10E-5), *CACNA1C* (cg19683021, P = 1.29E-6)*, CALN1* (cg14913143, P = 1.40E-6)*, DPYD* (cg21015639, P = 1.54E-5; cg24258243, P = 7.83E-5) and *GPR98* (cg15221739, P = 1.25E-5), and two SCHEMA genes (*TRIO* (cg12337605, P = 1.94E-6) and *CACNA1G* (cg11421573, p = 1.53E-5) (**Figure 2**). Overall, neuronal DMPs were significantly enriched at genes implicated by schizophrenia genetic studies (odds ratio = 3.24, P = 0.00380), demonstrating convergence between inherited genetic risk and schizophrenia-associated neuronal DNA methylation differences.

**Figure 2:**
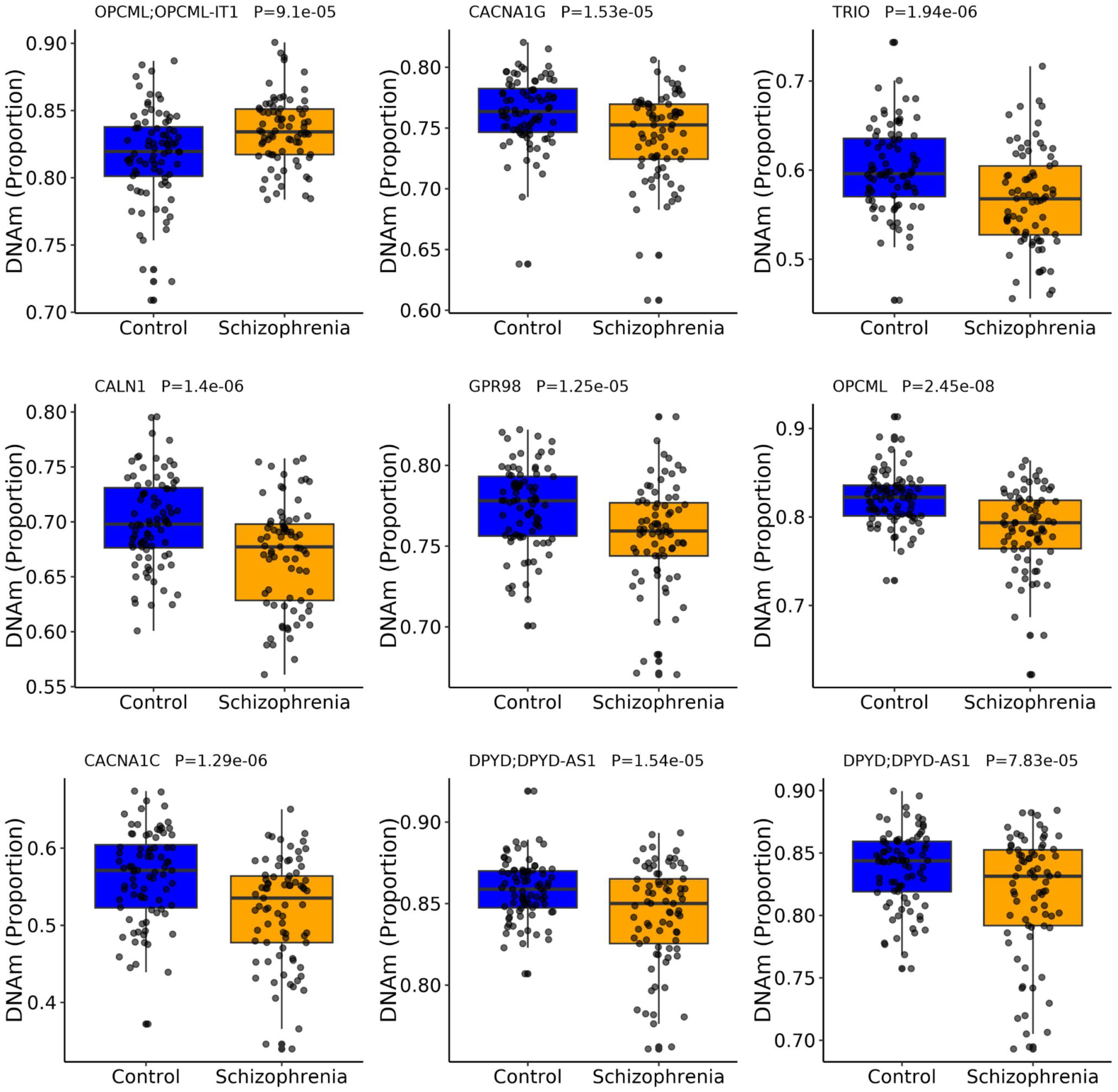
Convergence of schizophrenia-associated neuronal DNA methylation differences with genetic risk loci. Schizophrenia-associated neuronal differentially methylated positions (DMPs) overlapped genes implicated by both common and rare variant studies of schizophrenia. Nine DMPs mapped to five genes identified by the Psychiatric Genomics Consortium (PGC) (*OPCML*, *CACNA1C*, *CALN1*, *DPYD* and *GPR98*), while two DMPs mapped to genes implicated by rare coding variant analyses from the SCHEMA consortium (*TRIO* and *CACNA1G*).

To further investigate the biological processes implicated by schizophrenia-associated DNA methylation differences in neurons, we performed pathway enrichment analyses using both Gene Ontology Biological Process (GO-BP) and KEGG annotations, applying methods specifically adapted for DNA methylation data that account for probe number bias and inter-probe correlation^18^. These analyses identified highly significant enrichment of pathways involved in neuronal development, connectivity and synaptic function (**Figure 3** and **Supplementary Table 6**). Among the strongest GO-BP enrichments were terms related to homophilic cell adhesion via plasma membrane adhesion molecules (35 genes; FDR = 2.85E-26), nervous system development (113 genes; FDR = 4.54E-23), cell-cell adhesion via plasma membrane adhesion molecules (37 genes; FDR = 4.95E-21), and actin cytoskeleton organisation (42 genes; FDR = 4.50E-11). Notably, several highly enriched processes were directly related to synaptic biology, including synapse organisation (32 genes; FDR = 1.24E-7, chemical synaptic transmission (37 genes; FDR = 3.35E-7), anterograde trans-synaptic signalling (37 genes; FDR = 3.35E-7) and vesicle-mediated transport (56 genes; FDR = 3.36E-7). Consistent with these findings, KEGG pathway analysis highlighted significant enrichment of pathways involved in neuronal connectivity and neurotransmission, including cadherin signalling (37 genes; FDR = 1.39E-16), axon guidance (14 genes; FDR = 4.62E-5), immunoglobulin superfamily cell adhesion molecule signalling (18 genes; FDR = 4.62E-5), regulation of the actin cytoskeleton (15 genes; FDR = 1.17E-4) and the synaptic vesicle cycle (7 genes; FDR = 6.30E-3). Together, these analyses demonstrate that schizophrenia-associated neuronal DNA methylation differences are enriched in pathways related to neuronal adhesion, synapse formation, neurotransmission and activity-dependent plasticity, consistent with convergent genetic and functional evidence implicating disrupted cortical circuitry in schizophrenia^19^.

**Figure 3.**
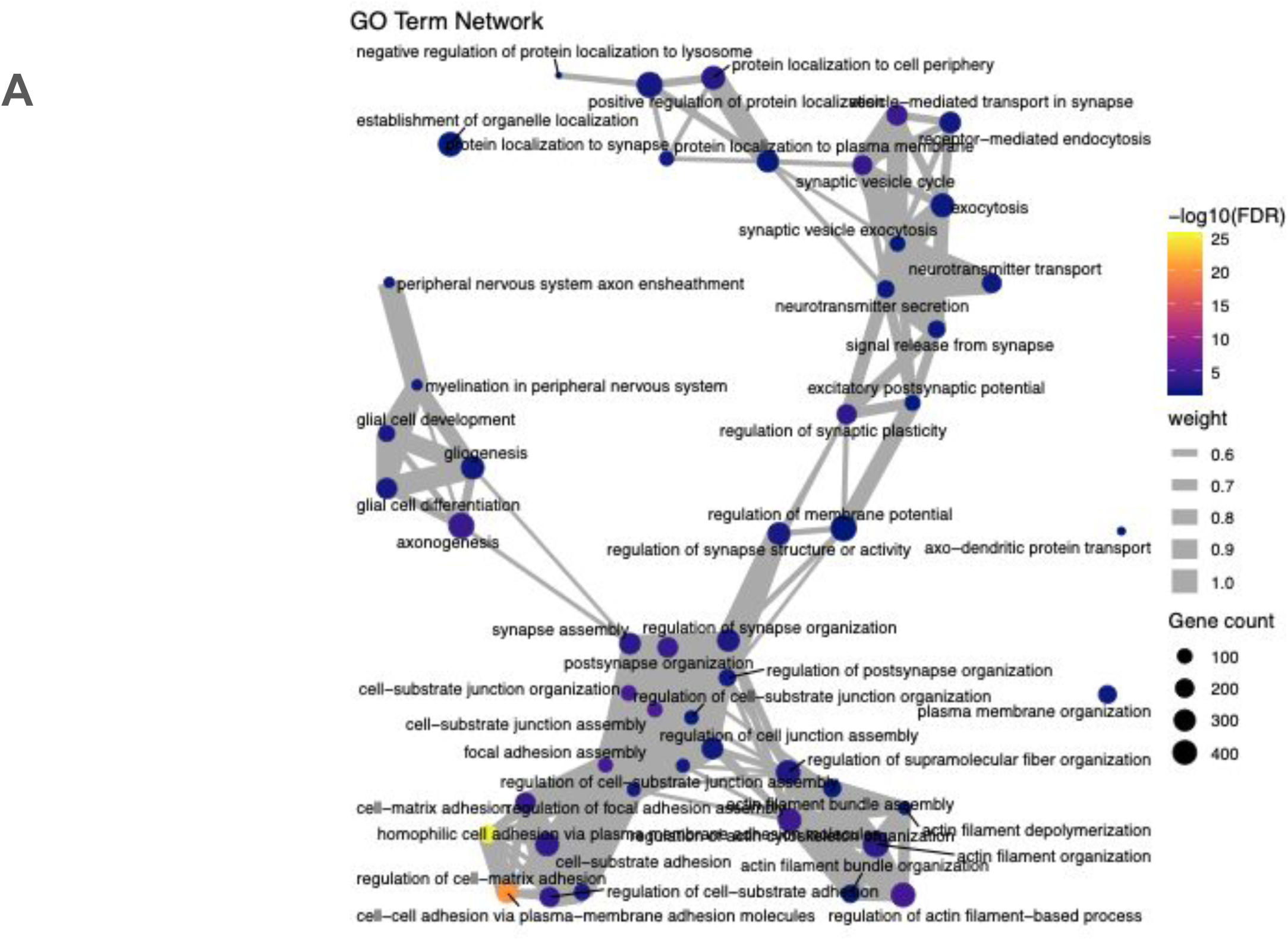
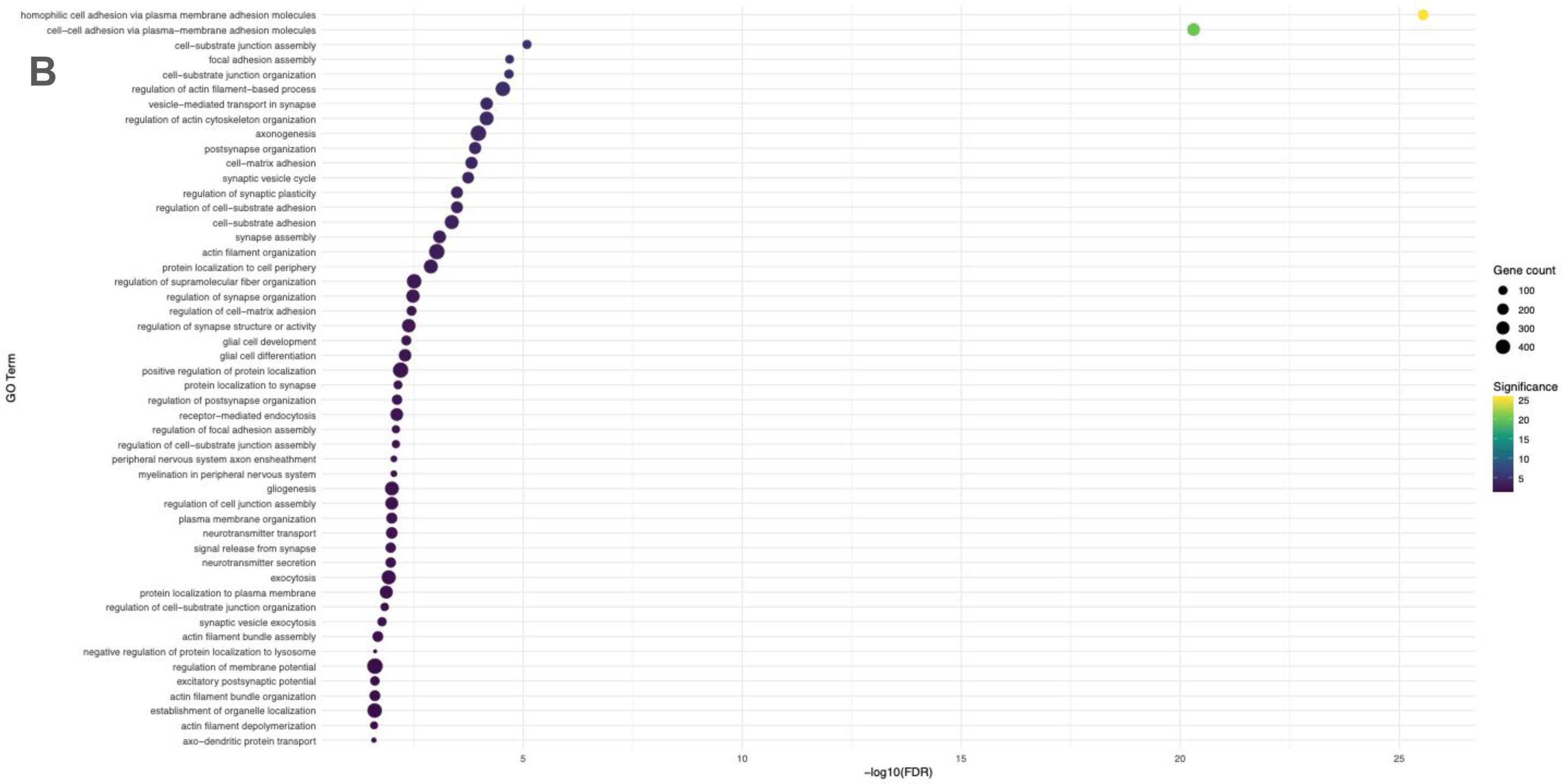
Gene Ontology enrichment analysis of schizophrenia-associated neuronal DNA methylation differences. **A)** Network plot showing relationships among the top 50 significantly enriched Gene Ontology Biological Process (GO-BP) terms identified from genes annotated to schizophrenia-associated differentially methylated positions in NeuN+ nuclei. Nodes represent individual GO-BP terms, with node size proportional to the number of genes within each term. Edges indicate overlap in gene membership between pathways, highlighting clusters of related biological processes. **B)** Bubble plot of the top 50 most significantly enriched Gene Ontology (GO) categories ranked by enrichment significance. Bubble size reflects the number of genes associated with each term, while colour indicates the strength of enrichment significance (−log10 adjusted P-value).

To obtain a more refined understanding of the synaptic processes implicated by schizophrenia-associated neuronal DMPs, we performed enrichment analyses using SynGO^20^, a curated expert-annotated knowledgebase of synaptic genes, components and biological processes (**Supplementary Table 7**). Consistent with the GO and KEGG analyses, schizophrenia-associated neuronal DMPs showed strong enrichment within synaptic compartments, including the synapse (97 genes; FDR = 6.20E-22), postsynapse (68 genes; FDR = 5.22E-17) and presynapse (48 genes; FDR = 1.24 E-10). Significant enrichment was also observed for specialised structures including the presynaptic active zone and postsynaptic actin cytoskeleton. Analysis of SynGO biological process annotations highlighted pathways involved in synapse organisation (38 genes; FDR = 5.30E-13), postsynaptic actin cytoskeleton organisation (7 genes; FDR = 2.50E-6), synaptic vesicle endocytosis (7 genes; FDR = 3.77E-3) and synapse maturation (5 genes; FDR = 4.35E-2). Together, these findings demonstrate that schizophrenia-associated PFC neuronal methylation differences preferentially affect genes involved in the formation, maintenance and activity-dependent remodelling of synapses, with particularly strong enrichment for pathways governing synaptic organisation and structural plasticity.

### Schizophrenia-associated neuronal DNA methylation signatures are partially conserved across brain regions

We next examined whether schizophrenia-associated DNA methylation differences identified in PFC neurons were also evident in neuronal nuclei isolated from two additional brain regions strongly implicated in schizophrenia, the hippocampus and striatum^21^. Using FANS we isolated NeuN+ nuclei from both brain regions for a subset of donors (n = 20, **Supplementary Table 1**) and profiled DNA methylation using the Illumina EPIC array as previously described (see **Methods**). As an initial validation, we confirmed that schizophrenia-associated effect sizes observed in PFC neurons from this subset closely recapitulated those identified in the full cohort. For both experiment-wide significant and discovery-threshold DMPs, direction of effect was highly concordant (100% and 99.3%, respectively), with strong correlations in effect size estimates (r = 0.87 and r = 0.95, respectively) (**Supplementary Figure 12**), indicating that the selected subset was highly representative of the larger study population. We then compared schizophrenia-associated effect sizes at PFC neuronal DMPs across hippocampal and striatal neurons. Overall, schizophrenia-associated methylation signatures were partially preserved across brain regions, with greater concordance observed in hippocampal neurons (stringent DMPs: concordant direction of effect = 92.9%, effect size corr = 0.42; discovery DMPs: concordance direction of effect = 74.9%, effect size corr = 0.41) than striatal neurons (stringent DMPs: concordant direction of effect = 85.7%, effect size corr = 0.12; discovery DMPs: concordance direction of effect = 55.4%, effect size corr = 0.20) (**Supplementary Figure 12**). These findings suggest that a subset of schizophrenia-associated neuronal methylation differences are consistent across multiple brain regions, while others exhibit greater regional specificity. This heterogeneity was evident at individual loci. Although some PFC NeuN+ DMPs were not consistent across brain regions (e.g. hypomethylation at cg08080338 annotated to HUNK was restricted to PFC neurons (matched PFC NeuN+ nuclei P = 0.000816, hippocampus NeuN+ nuclei P = 0.417, striatum NeuN+ nuclei P = 0.615)), others were characterized by consistent differences in NeuN+ nuclei across brain regions (e.g. cg12421899 annotated to SS5CD showed consistent schizophrenia-associated DNA methylation differences in neuronal nuclei from all three brain regions (matched PFC NeuN+ nuclei P = 0.000278, hippocampus NeuN+ nuclei P = 0.0320, striatum NeuN+ nuclei P = 0.00413) (**Supplementary Figure 13**). These findings indicate that schizophrenia-associated neuronal epigenetic differences might reflect both region-specific and cross-regional components, potentially reflecting distinct aspects of disease pathology.

## DISCUSSION

This study represents, to our knowledge, the largest and most comprehensive cell type-resolved analysis of DNA methylation variation in schizophrenia to date. By profiling purified neuronal, oligodendrocyte and glial nuclei populations from human prefrontal cortex, we demonstrate that schizophrenia-associated methylomic variation is highly cell type-specific and overwhelmingly localised to neurons. All experiment-wide significant differentially methylated positions were identified exclusively in neuronal nuclei and were characterised by a pronounced bias towards hypomethylation, with no significant associations detected in oligodendrocyte-enriched, glial-enriched or bulk tissue fractions. These neuronal DNA methylation differences converged on genes and pathways implicated by genetic studies of schizophrenia, particularly those involved in synaptic organisation, neuronal connectivity and activity-dependent plasticity. Together, our findings support a model in which schizophrenia-associated molecular dysregulation is concentrated within neuronal regulatory networks and highlight the limitations of conventional bulk tissue approaches for investigating disease mechanisms in the human brain.

Previous studies have highlighted altered neuronal activity states and dysregulated activity-dependent gene regulation in schizophrenia^22,23^, and it is known that epigenetic regulation in neurons can be highly dynamic and responsive to synaptic activity^24^. We found an enrichment of neuronal DMPs within pathways related to neurotransmission, calcium signalling, synaptic organisation and cytoskeletal regulation, which is interesting given long-standing evidence implicating disrupted cortical circuitry and synaptic dysfunction in schizophrenia^4,25^. Importantly, schizophrenia-associated neuronal DMPs significantly overlapped loci implicated by large-scale genetic studies, including genes identified through common and rare variant analyses such as *CACNA1C*, *CACNA1G* and *TRIO*. This convergence suggests that at least a subset of the observed methylomic differences potentially reflect downstream regulatory consequences of schizophrenia genetic risk. Many schizophrenia risk variants reside within non-coding regulatory regions active in cortical neurons^26^, and our findings are consistent with the hypothesis that genetic risk influences disease through disruption of neuron-specific regulatory mechanisms. Whether the DNA methylation differences identified here represent causal mediators of genetic risk, compensatory responses, or secondary consequences of disease-related processes remains unclear, but the overlap with established schizophrenia loci strengthens their biological relevance.

One of the most important findings from this study is the extent to which bulk tissue analyses obscure schizophrenia-associated neuronal methylation differences. Although neuronal DMP effect sizes were directionally correlated with those observed in bulk cortex, effect sizes were substantially attenuated and no experiment-wide significant loci were identified in matched total nuclei fractions. These observations potentially explain the limited replication and heterogeneity that have characterised previous schizophrenia EWAS using bulk brain tissue. Cellular heterogeneity is a major confounder in epigenomic studies of complex tissues, particularly in psychiatric disorders, where subtle within-cell molecular alterations may be masked by inter-individual variation in cell composition, agonal state, medication exposure and tissue quality. Our results demonstrate that direct purification of disease-relevant cell populations substantially increases sensitivity for detecting biologically meaningful disease-associated variation.

Interestingly, schizophrenia-associated neuronal DNA methylation differences identified in PFC also showed correlated effects in neuron-enriched nuclei isolated from hippocampus and striatum. Despite the limited power of these analyses given the relatively small subset of samples assessed, these findings suggest that schizophrenia-associated neuronal methylation differences may potentially reflect broader circuit-level or brain-wide processes rather than being brain region-specific. Of note, however, the reduced concordance observed in striatal neurons also points towards regional specificity in the epigenetic architecture of schizophrenia, potentially reflecting differences in neuronal composition, developmental origin or circuit function across brain regions.

This study has several important limitations that should be considered when interpreting our findings. First, although substantially larger than previous cell-type-specific brain EWAS studies, our sample size remains modest, especially relative to genetic studies of schizophrenia. Although this may limit power to detect smaller effects, we have increased power relative to bulk tissue studies of the same size. Second, DNA methylation differences identified in post-mortem tissue cannot readily distinguish causal disease mechanisms from secondary effects related to chronic illness, medication exposure, smoking or other environmental factors associated with schizophrenia. Third, while our nuclei sorting strategy substantially improves cellular resolution, the neuronal and glial populations studied here still comprise heterogeneous cell subtypes. Emerging single-cell multi-omic technologies will ultimately be required to define schizophrenia-associated epigenetic alterations at finer cellular resolution. Fourth, while the Illumina EPIC array enables precise quantification of DNA methylation at single-base resolution for sites annotated to the vast majority of genes and is more comprehensive than assays used in previous studies of schizophrenia, it covers only a small proportion of CpG sites in the human genome. As sequencing costs decline, future research should utilize sequencing-based technologies to comprehensively profile DNA methylation differences in schizophrenia. Fifth, our study could not distinguish between DNA methylation and its oxidized form, DNA hydroxymethylation, due to limitations of sodium bisulfite-based approaches^27^. This distinction is potentially important, as DNA hydroxymethylation is abundant in the central nervous system and known to impact gene expression^28,29^. Future studies should aim to differentiate these modifications using techniques such as nanopore sequencing^30^. Lastly, we did not obtain gene expression data from these samples, preventing direct conclusions about the transcriptional impact of the observed DNA methylation changes. Integration with chromatin accessibility, histone modification and transcriptomic datasets will be necessary to fully understand the functional consequences of the DNA methylation differences identified here.

In summary, this study demonstrates that schizophrenia-associated cortical DNA methylation variation is highly neuron-specific, characterised by widespread hypomethylation and concentrated within genes and pathways governing synaptic organisation, neuronal connectivity and schizophrenia genetic risk. By revealing disease-associated methylation differences that are largely obscured in bulk tissue analyses, our findings redefine the cellular landscape of epigenetic dysregulation in schizophrenia. These results provide a framework for integrating genetic, epigenomic and functional genomic data to elucidate how risk for schizophrenia converges on neuronal regulatory networks and ultimately disrupts cortical circuit function.

## METHODS

### Isolation of cell-type-specific nuclei populations from post-mortem cortex tissue

Post-mortem cortex tissue was provided from multiple brain banks from the UK, Canada and USA. Subjects were approached in life for written consent for brain banking, and all tissue donations were collected and stored following legal and ethical guidelines. All samples were dissected by trained neuropathologists from each brain bank, snap-frozen and stored at −80°C before being transferred to the University of Exeter under Material Transfer Agreements. An overview of the samples used in this study is provided in **Supplementary Table 1**.

### Fluorescence-activated nuclei sorting of different nuclei populations from the PFC

∼500mg PFC tissue was processed using a FANS protocol developed by our group^14^. A full protocol detailing each step of our nuclei purification protocol is also provided on protocols.io at https://dx.doi.org/10.17504/protocols.io.36wgq4965vk5/v3. Briefly, following tissue homogenization and nuclei purification using sucrose gradient centrifugation we used a FACS Aria III cell sorter (BD Biosciences) to simultaneously collect populations of NeuN+ (neuronal-enriched) (R&D systems, Cat No: NL2864R, dilution: 1:10) and SOX10+ (oligodendrocyte-enriched) (Millipore, Cat No: MAB377X, dilution: 1:1000) immunolabeled populations from bulk DLPFC tissue prior to genomic profiling, with the double-negative fraction (other glia-enriched) and an aliquot of the ‘total’ nuclei fraction (analogous to bulk cortex) also being collected from each tissue sample. Nuclei suspensions were assessed for the presence of debris by adjusting the gating strategy before proceeding with nuclei capture. For each sorted population, ∼200,000 nuclei were collected for extraction of genomic DNA using a phenol:chloroform extraction protocol optimised for FANS-isolated nuclei^31^. Details of the gating strategies we implemented are shown in **Supplementary Figure 1**.

### DNAm profiling, processing and quality control

500 ng of genomic DNA from each sample was treated with sodium bisulfite using the Zymo EZ-96 DNA Methylation-Gold™ Kit (Cambridge Bioscience, UK) according to the manufacturer’s standard protocol. All samples were then processed using the Illumina EPIC HumanMethylation 850K array (Illumina Inc, CA, USA) according to the manufacturer’s instructions, with minor amendments and quantified using an Illumina iScan System (Illumina, CA, USA). Individuals were randomised and sorted fractions from the same individual and FANs gating run were processed on the same BeadChip, where within a BeadChip the location of each fraction was randomised. DNAm data was loaded into R (version 3.6.3) from idat files using the package *bigmelon*^32^. These data were processed through a bespoke quality control pipeline developed for cell-specific DNAm data^33^. The final dataset comprised of 653 nuclei fractions (NeuN+: cases = 77, controls = 84; SOX10+: cases = 69, controls = 84; NeuN-/SOX10-: cases = 65, controls = 77; total nuclei: cases = 96, controls = 101). Data were normalised with the *dasen*() function in the *wateRmelon* R package^34^ separately for each cell type. The composition of each nuclei fraction was estimated for each sample using brain-specific reference panels provided through the *CETYGO* R package^16^ and we used a t-test to assess differences in sorting efficiency between cases and controls.

### Single nucleus RNA-seq

To further validate the purity of the sorted nuclei populations, we performed single-nucleus RNA sequencing (snRNA-seq) on PFC nuclei isolated from four donors. Minor modifications were made to the FANS protocol to optimise RNA integrity, including the addition of Ribolock RNase inhibitor (Thermo Scientific, EO0382; 0.2 U/ml) to both the lysis and staining buffers. For each nuclei fraction, 100,000 sorted nuclei were fixed using the Parse Evercode Low-Input Nuclei Fixation Kit (v3), with approximately 6,250 nuclei targeted for recovery per sample. snRNA-seq libraries were prepared using the Evercode WT Kit (v3) and library quality assessed using the Agilent D5000 High Sensitivity ScreenTape assay. Libraries were pooled and sequenced on an Illumina NovaSeq 6000 platform (100 bp paired-end reads) to a target depth of ∼20,000 reads per nucleus. Sequencing reads were aligned to the GRCh38 reference genome and filtered count matrices generated using Trailmaker (v1.5.1, Parse Biosciences), excluding barcodes containing fewer than 10 transcripts or failing the barcode rank plot threshold. Putative doublets were identified and removed using scDblFinder^35^.

Downstream analyses were performed in Seurat (v5.2.1)^36^. Nuclei with at least 200 detected transcripts and fewer than 10% mitochondrial reads were retained, alongside genes detected in at least five nuclei. Data from each sample were log-normalised and scaled, with the top 4,000 variable features selected for dimensionality reduction by principal component analysis (PCA). To account for inter-sample technical variation, datasets were integrated using Harmony^37^ prior to graph-based clustering using the Leiden community detection algorithm. Uniform Manifold Approximation and Projection (UMAP) embeddings were generated for the visualisation of cellular structure. Following a second round of filtering excluding nuclei with fewer than 400 detected genes, clusters were manually annotated as major neural cell populations based on established marker gene expression profiles.

### Cell specific Epigenome-wide Association Analysis

Sites previously identified as potentially cross-hybridising, containing a common polymorphism in the hybridisation sequence (> 1% minor allele frequency)^34,38^, flagged by the manufacturer as affected by variable technical performance or on the Y chromosome, were excluded from analysis. To identify sites where DNA methylation level differed between cases and controls a linear regression model was fitted within each cell type. For each cell type and each DNA methylation site, DNA methylation level was treated as the dependent variable with a binary variable for case-control status, and covariates for age, sex, and brain bank. For the regression analysis with NeuN+ samples, we additionally fitted a second model that controlled for the proportion of SOX6+ nuclei. For all regression models, DMPs were identified from the significance testing of the case control difference using Student’s t-distribution and using our previously established experiment-wide significance threshold (P < 9.00E-8) and a discovery threshold of P < 1.00E-4. The second stage of the analysis was to confirm whether these were specific to the cell type in which they were discovered. For this analysis, we used all available samples across all fractions. We fitted a mixed-effects model, with DNA methylation level as the dependent variable, case-control status, age, sex, and cell type as fixed effects, and individual and tissue centre as random effects. In addition, to allow for differences in the schizophrenia associated effect between cell types, an interaction term was included between case-control status and cell type. These models were fitted with the R package *lme4* and significance values calculated using the R package *lmerTest*. An ANOVA was used to test for significant differences by comparing the full model with the interaction term to a null model without it.

### Gene ontology pathway analysis

Pathway analysis was performed using the *gsGene* function from the *dmGsea* package in R^18^. Briefly, probe-level P values were combined into gene-level P values using Fisher’s method with Nyholt correction for probe correlation. Gene set enrichment was then tested against the KEGG and GO-BP databases using the threshold method (FDR < 0.05). To obtain a more refined understanding of synaptic processes implicated by schizophrenia-associated neuronal DMPs, analyses were also performed using SynGO^20^, a curated expert-annotated knowledgebase of synaptic genes, components and biological processes.

### Bisulfite pyrosequencing

Neuron-enriched nuclei populations (50,000 nuclei per sample) were subjected to direct bisulfite conversion using the Zymo Research Direct Bisulfite Kit prior to PCR amplification of two regions within *HUNK* using primers designed against bisulfite-converted DNA. For assay 1 (targeting CpG3 and CpG4), the forward primer sequence was 5′-GGGGAGTGGTTTTTGTATTAAGTTATGA-3′, the biotinylated reverse primer was 5′-CATACATTCTATCACTCTCCAAATACT-3′, and the sequencing primer was 5′-GTAGATGTTGTTGTATAGG-3′. For assay 2 (targeting CpG1 and CpG2), the biotinylated forward primer was 5′-TGGTGATTTTATTGGTTAGTTATTATTAGT-3′, the reverse primer was 5′-CCCTATACAACAACATCTACAATAAAATAT-3′, and the sequencing primer was 5′-ACAAAACCACTCC-3′. PCR amplification was performed under the following cycling conditions: initial enzyme activation at 95°C for 10 min, followed by 38 cycles of denaturation at 94°C for 30 s, annealing at 50°C (assay 1) or 60°C (assay 2) for 1 min, and extension at 72°C for 1 min, with a final extension step at 72°C for 10 min. Pyrosequencing was performed on the PyroMark Q48 platform (Qiagen, UK). Biotin-labelled PCR products were immobilised on streptavidin-coated beads and denatured to generate single-stranded DNA templates prior to sequencing primer annealing. DNA methylation levels at individual CpG sites were quantified from the relative incorporation of cytosine and thymine nucleotides during pyrosequencing^39^.

## ACKNOWLEDGEMENTS

This project was funded by Medical Research Council (MRC) grants K013807 and W004984 (awarded to J.M.). E.H is supported by an Engineering and Physical Sciences Research Council Fellowship EP/V052527/1. Data analysis was undertaken using high-performance computing supported by an MRC Clinical Infrastructure award (M008924) to J.M. Method development was supported by Alzheimer’s Research UK (ARUK) grant ARUK-PPG2018A-010 to E.L.D. This study was also supported by the National Institute for Health and Care Research Exeter Biomedical Research Centre. The views expressed are those of the author(s) and not necessarily those of the NIHR or the Department of Health and Social Care. We acknowledge the supply of samples from: The Cambridge Brain Bank, covered by current REC approval NRES 10/HO308/56; the Quebec Suicide Brain Bank at Douglas Mental Health University Institute, Canada; the MRC funded University of Edinburgh Brain & Tissue Bank; the Harvard Brain Tissue Resource Centre, which is supported by HHSN-271-2013-00030C; Brain Endowment Bank, at Miller School of Medicine, University of Miami; The Mount Sinai NBTR (NIH Brain and Tissue Repository), JJ Peters VA Medical Center; the Oxford Brain Bank, supported by the Medical Research Council (MRC), the NIHR Oxford Biomedical Research Centre and the Brains for Dementia Research programme, jointly funded by Alzheimer’s Research UK and Alzheimer’s Society; The Neuropathology Brain Bank at the University of Pittsburgh, School of Medicine Department of Psychiatry; The Stanley Medical Research Institute Brain Collection courtesy of Drs. Michael B. Knable, E. Fuller Torrey, Maree J. Webster, and Robert H. Yolken; The Human Brain and Spinal Fluid Resource Center (HBSFRC) NIH Neurobiobank.

## Notes

### Competing Interest Statement

The authors have declared no competing interest.

