## Supplementary Figures for "Schizophrenia-associated DNA methylation differences in the cortex are neuron-specific"

**Supplementary Figure 1: Fluorescence-activated nuclei sorting (FANS) workflow for generating cell type-specific DNA methylation data from human prefrontal cortex.** Full experimental details are provided in Chioza *et al.* (2025) and in the accompanying protocol available at https://www.protocols.io/view/fluorescence-activated-nuclei-sorting-fans-on-huma-36wgq4965vk5/v2. **A)** Schematic overview of the experimental workflow used to generate genome-wide DNA methylation profiles from purified nuclei populations isolated from post-mortem prefrontal cortex (PFC) tissue. Bulk cortical tissue was homogenised to release nuclei, followed by fluorescence-based immunolabelling, FANS purification of discrete nuclei populations, DNA extraction and Illumina EPIC array profiling. **B)** Representative FANS gating strategy used to isolate major cortical nuclei populations. Sequential gating steps were applied to remove debris, doublets and non-nuclear events prior to separation of nuclei populations according to NeuN–Alexa Fluor 488 and SOX10–NL577 fluorescence intensity. The scatterplot illustrates the identification of NeuN+ neuronal nuclei (purple) and SOX10+ oligodendrocyte nuclei (dark pink), with NeuN−/SOX10− nuclei representing the remaining glial-enriched population.

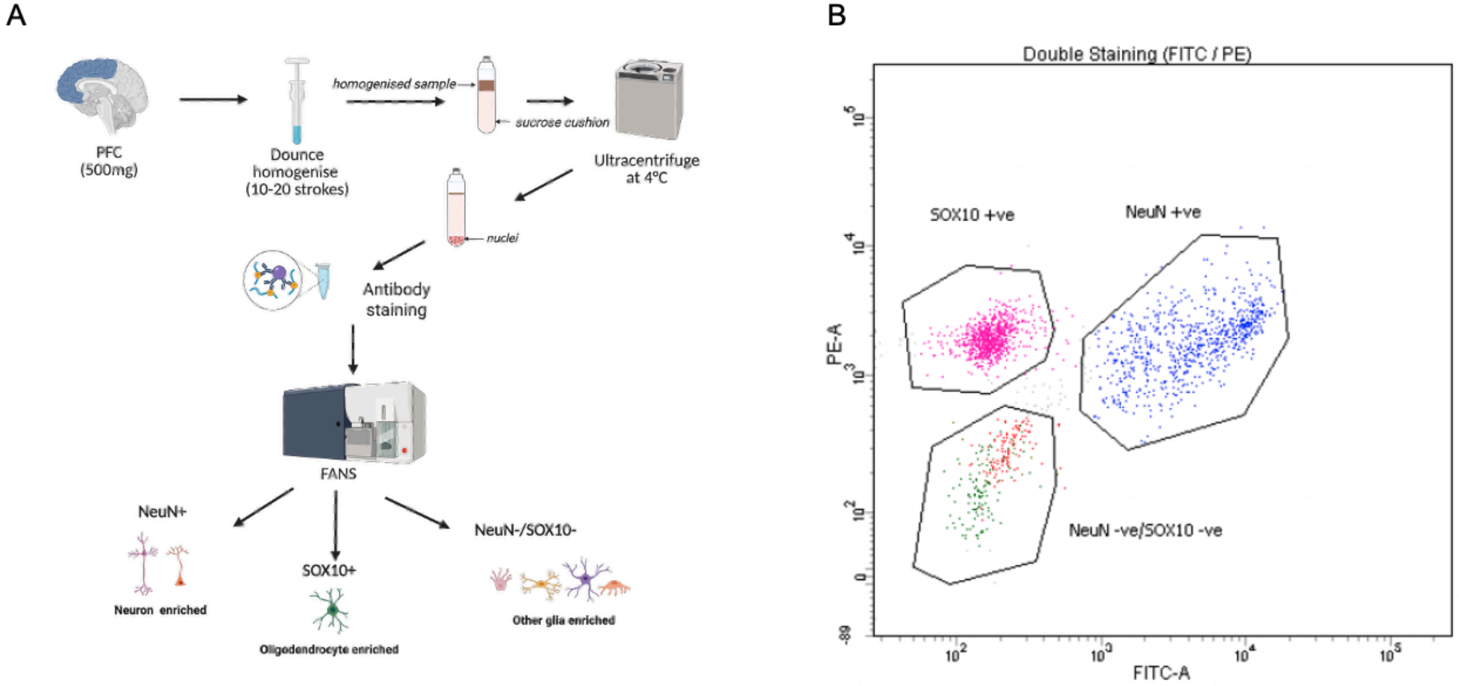

**Supplementary Figure 2: Single-nucleus RNA sequencing confirms efficient purification of cell type-specific nuclei populations by fluorescence-activated nuclei sorting.** Three FANS-isolated nuclei fractions from four post-mortem PFC samples were profiled using the Parse Biosciences Evercode low-input fixation workflow followed by single-nucleus RNA sequencing. **A)** Integrated UMAP embedding showing nuclei coloured according to their originating FANS-isolated population (NeuN+, SOX10+ and NeuN−/SOX10− fractions). **B)** UMAP embedding coloured by inferred cell type identity following unsupervised clustering and annotation using the Seurat analysis workflow. Clusters were annotated on the basis of established neural cell type marker genes. **C)** Mean expression of RBFOX3 (encoding NeuN, left panel) and SOX10 (right panel) across the three FANS-isolated nuclei populations. Expression values are shown as log1p-transformed counts per 100,000 transcripts.

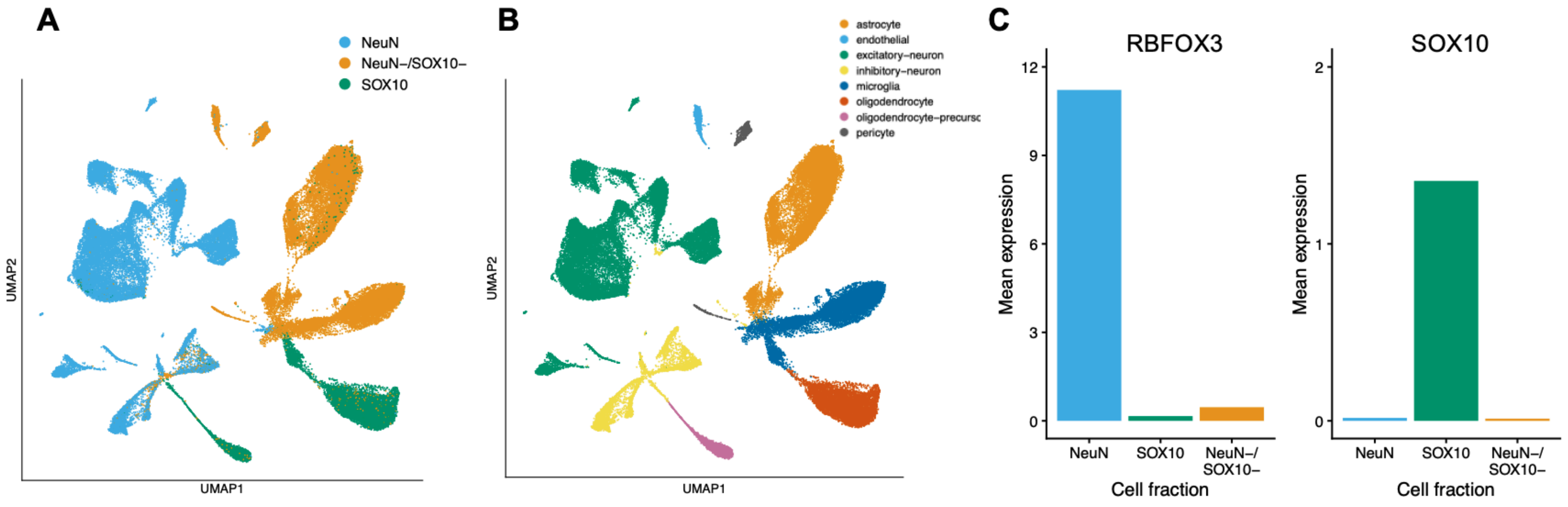

**Supplementary Figure 3: Validation of FANS purification efficiency using the *CETYGO* cellular deconvolution algorithm.** The composition of each nuclei fraction was estimated for each sample using brain-specific reference panels provided through the *CETYGO* R package (see **Methods**).

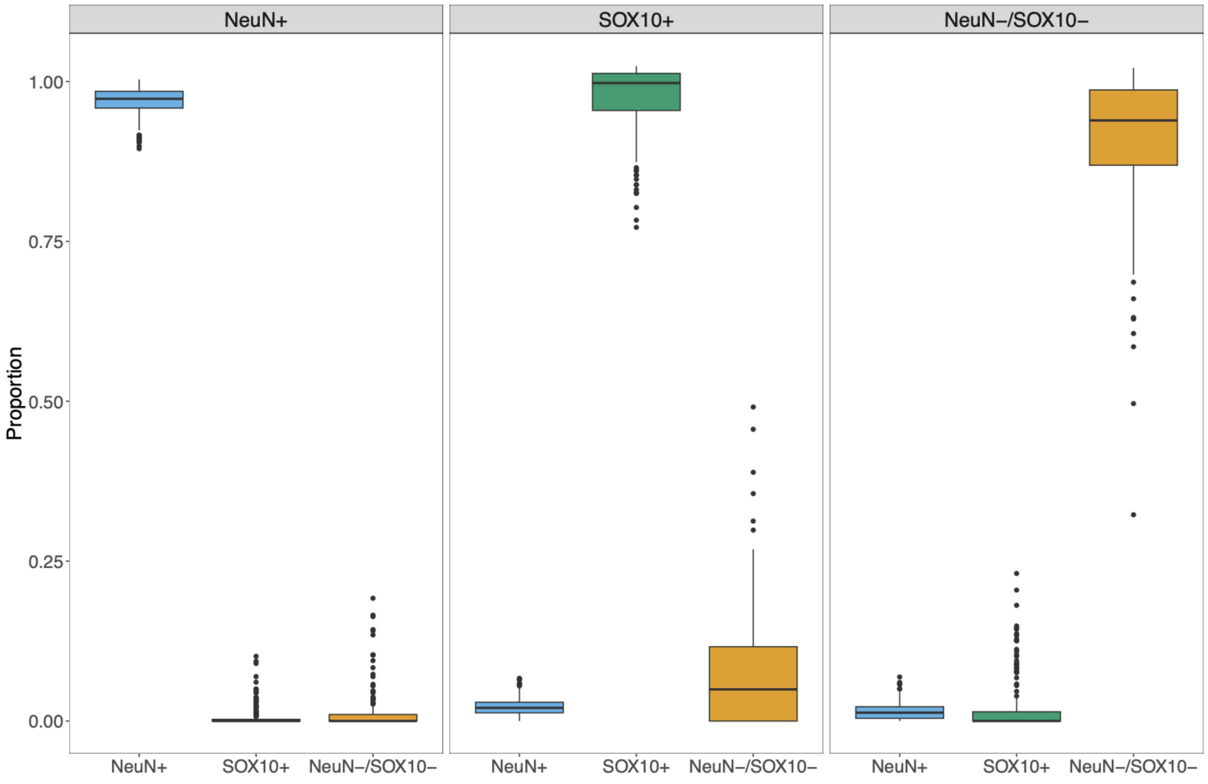

**Supplementary Figure 4: Unsupervised analyses demonstrate distinct DNA methylation signatures across purified cortical nuclei populations. A)** Principal component analysis (PCA) of genome-wide DNA methylation data from purified neuronal (NeuN+), oligodendrocyte (SOX10+) and other glial (NeuN−/SOX10−) nuclei fractions isolated from prefrontal cortex tissue. Samples cluster according to nuclei population, reflecting strong cell type-specific differences in cortical DNA methylation profiles. **B)** Hierarchical clustering heatmap generated using the 10,000 most variably methylated CpG sites across all samples. DNA methylation levels are represented from low methylation (blue; 0%) to high methylation (red; 100%). Clustering patterns further confirm the marked epigenetic separation of purified nuclei populations according to cell type identity.

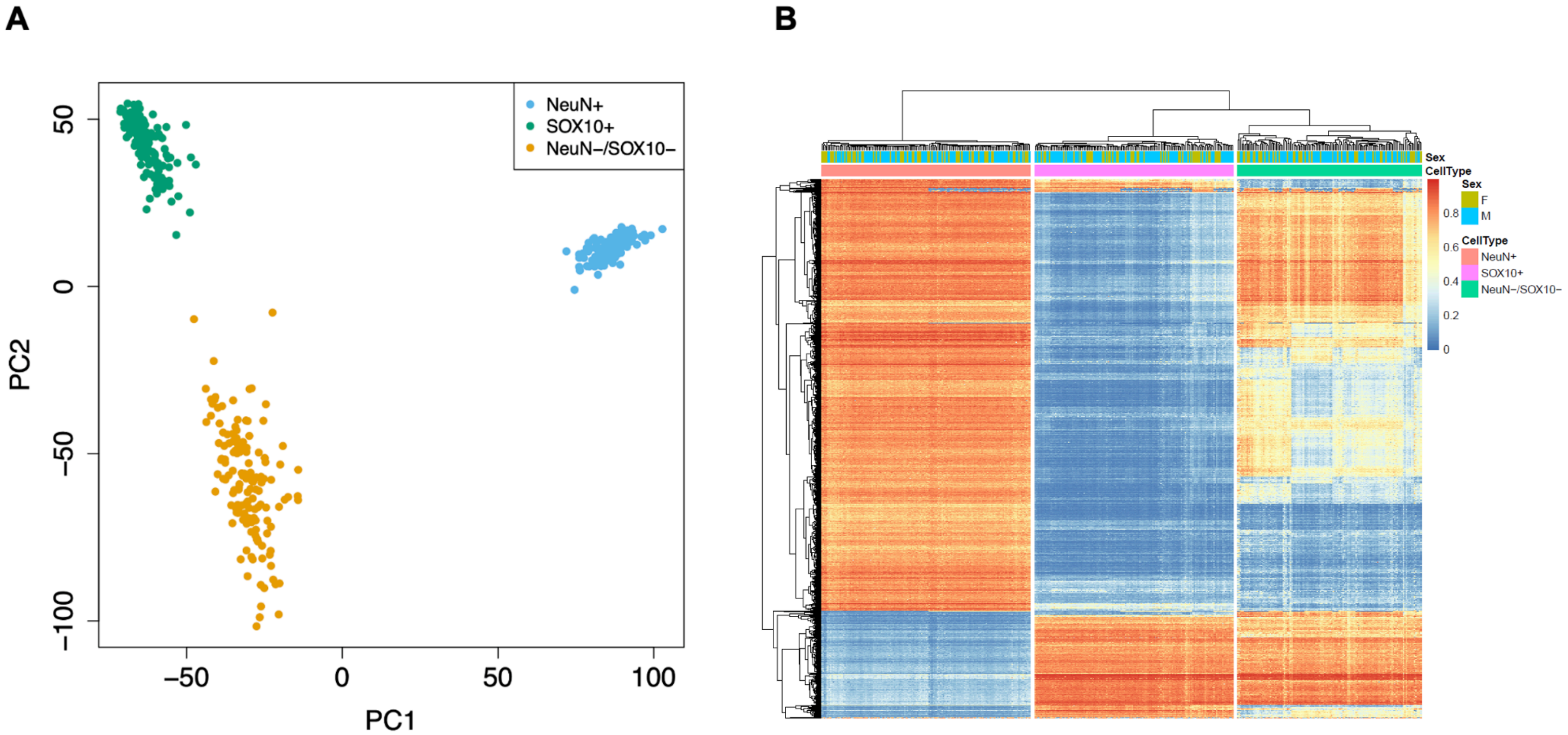

**Supplementary Figure 5: Quantile-quantile (QQ) plots indicate minimal genomic inflation in schizophrenia EWAS analyses across purified nuclei populations.** QQ plots comparing observed versus expected distributions of association test statistics for epigenome-wide association studies (EWAS) performed separately within neuronal (NeuN+), oligodendrocyte (SOX10+), other glial (NeuN−/SOX10−) and total nuclei fractions. Across all four analyses, observed P-value distributions closely matched the null expectation except for deviation in the extreme tail in NeuN+ nuclei, consistent with the presence of true schizophrenia-associated signals rather than widespread inflation driven by technical or confounding factors. Genomic inflation statistics (lambda) for each analysis are provided.

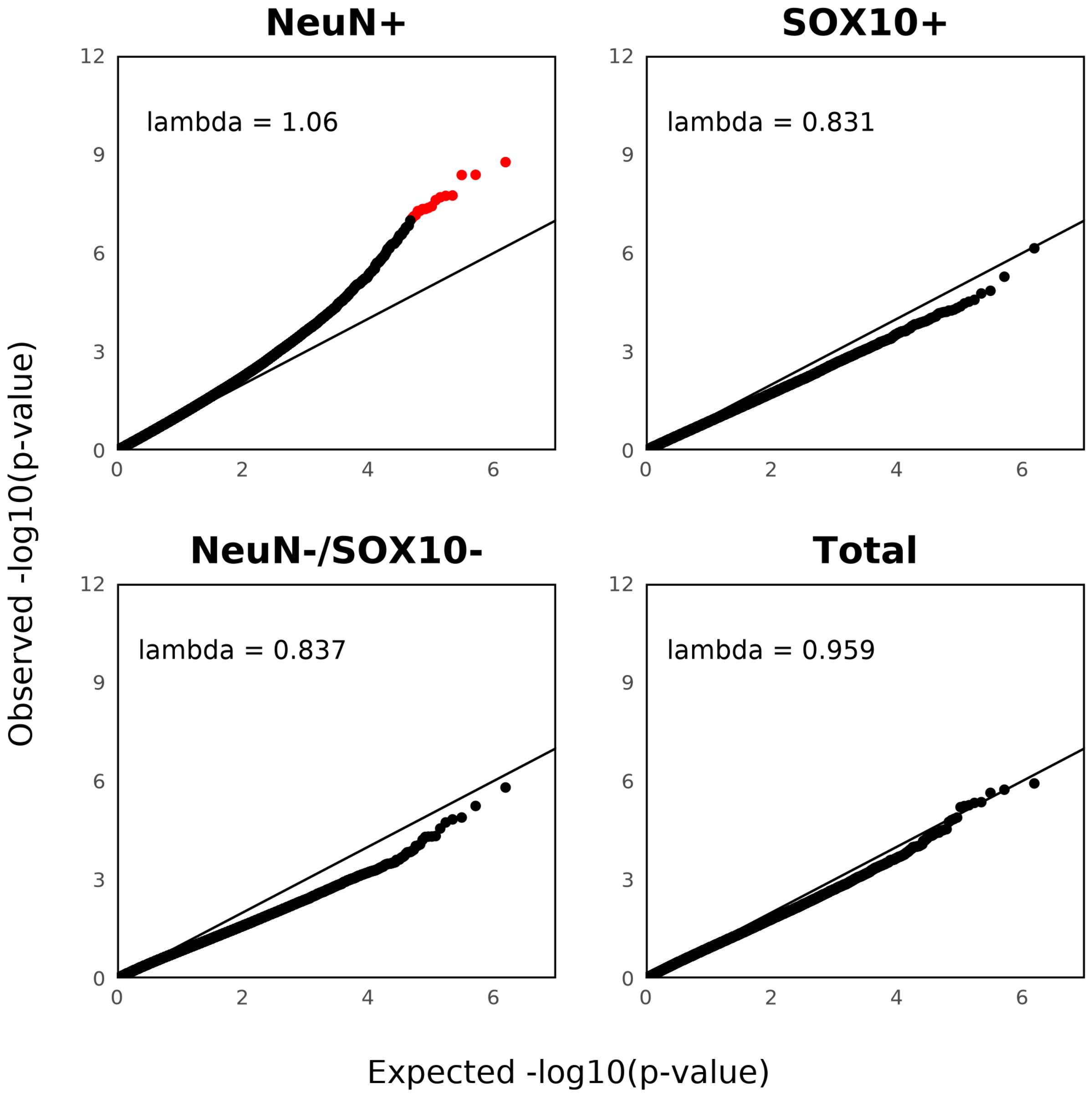

**Supplementary Figure 6: Correlation of effect sizes across different enriched nuclei fractions for DMPs identified in neuron-enriched (NeuN+) nuclei.** Scatterplots showing the correlation of schizophrenia EWAS effect sizes between neuron-enriched (NeuN+) nuclei and the other nuclei fractions (SOX10+, NeuN−/SOX10− and total nuclei fractions) for the 426 neuronal DMPs identified at the discovery significance threshold ($P < 1.00\times10^{-4}$). Although schizophrenia-associated effect estimates in NeuN+ nuclei were directionally correlated with those observed in the total nuclei fraction (corr = 0.54), effect sizes were markedly attenuated highlighting the increased sensitivity of cell type-resolved profiling approaches (**Figure 1B**). Each dot represents a DMP, with those reaching experiment-wide significance in neuronal nuclei ($P < 9.00\times10^{-8}$) shown in red.

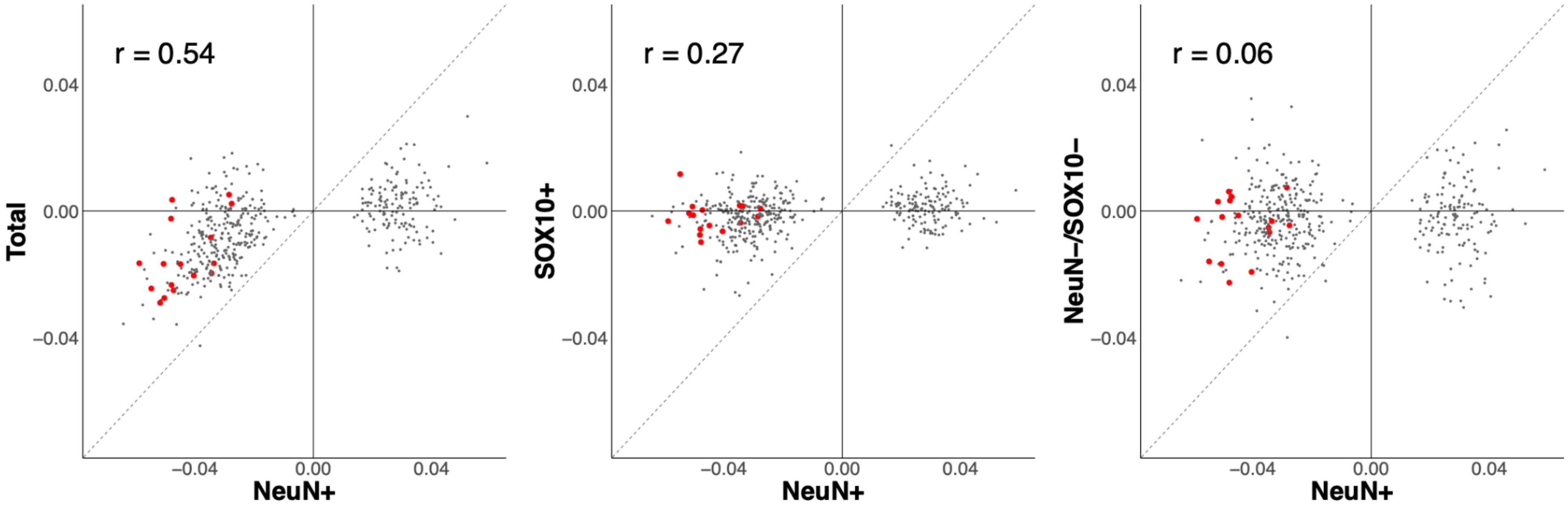

**Supplementary Figure 7: Schizophrenia-associated neuronal DMPs are unaffected by differences in inhibitory neuronal composition.** Scatter plot comparing schizophrenia-associated effect size estimates for discovery-threshold neuronal DMPs obtained from EWAS models with and without adjustment for the estimated proportion of SOX6+ inhibitory neurons within the NeuN+ fraction. SOX6+ neuronal abundance was estimated using CETYGO and reference DNA methylation data generated from purified SOX6+ nuclei. Effect size estimates were highly concordant between models (r = 0.996), demonstrating that the identified neuronal methylation differences are robust to variation in inhibitory neuronal representation. Each point represents a single neuronal DMP and the dashed line denotes the line of identity.

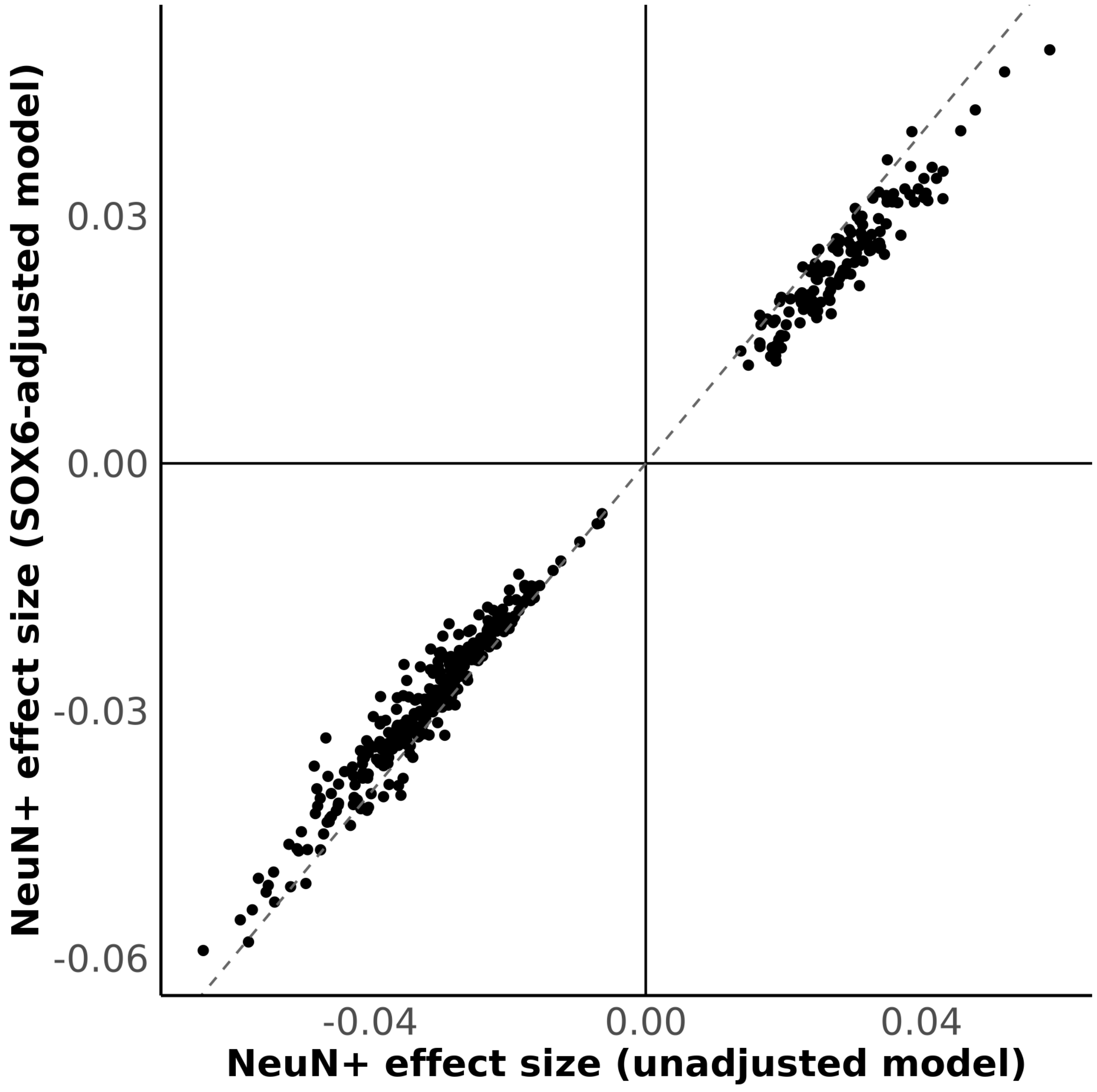

**Supplementary Figure 8: Genomic localisation of the schizophrenia-associated DMP cg08080338 within *HUNK*.** Mini-Manhattan plot showing association statistics across the HUNK locus together with gene annotations, EPIC array probe locations and ENCODE neuronal enhancer annotations.

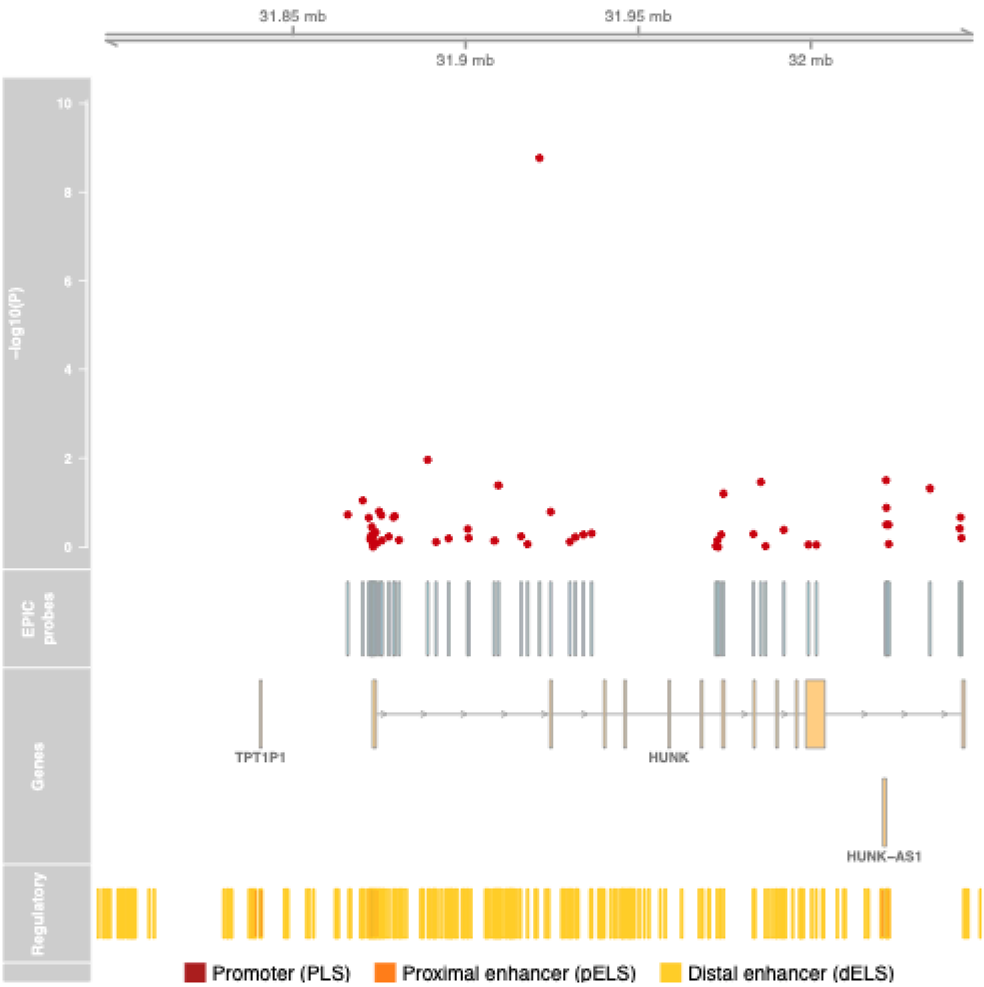

**Supplementary Figure 9: Schizophrenia-associated hypomethylation at cg08080338 is specific to neuronal nuclei.** Boxplots showing DNA methylation levels at cg08080338 (annotated to *HUNK*), the most significant schizophrenia-associated differentially methylated position (DMP) identified in neuronal nuclei, across schizophrenia cases and controls within each purified nuclei population. Significant hypomethylation was observed specifically in neuron-enriched (NeuN+) nuclei from schizophrenia cases, whereas no significant case-control differences were detected in oligodendrocyte-enriched (SOX10+) or other glial (NeuN−/SOX10−) nuclei populations.

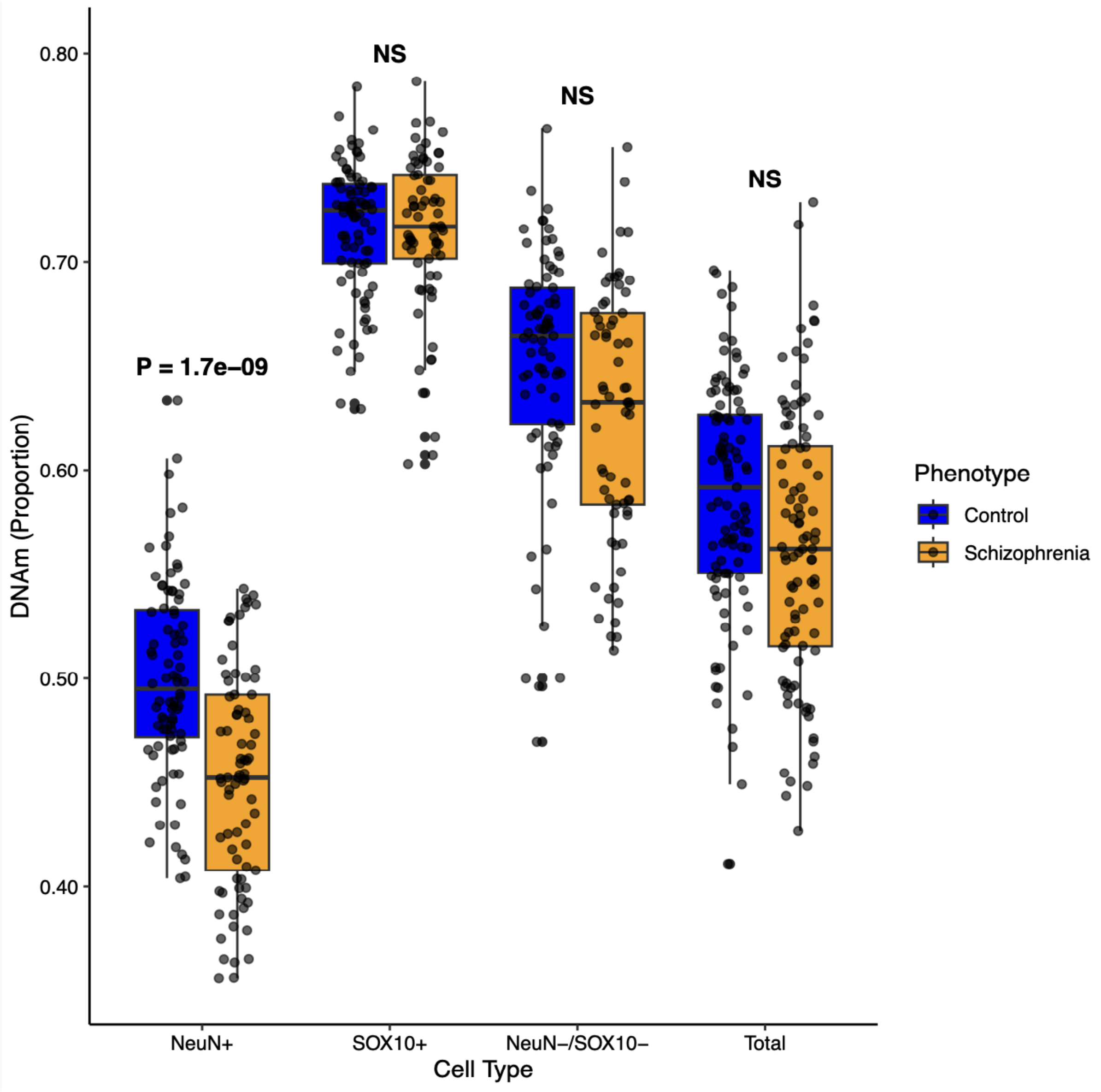

**Supplementary Figure 10: Bisulfite pyrosequencing confirms schizophrenia-associated hypomethylation at sites annotated to *HUNK* in neuron-enriched nuclei.** Bisulfite pyrosequencing was used to quantify DNA methylation across four CpG sites within targeted *HUNK* amplicons in NeuN+ (neuron-enriched) nuclei isolated by FANS from a subset of samples (n = 32). This included one site corresponding to cg08080338 (CpG2 in the assay) and three adjacent CpGs. Schizophrenia-associated hypomethylation was consistently observed at each site and across the broader amplicon region ($P < 0.03$).

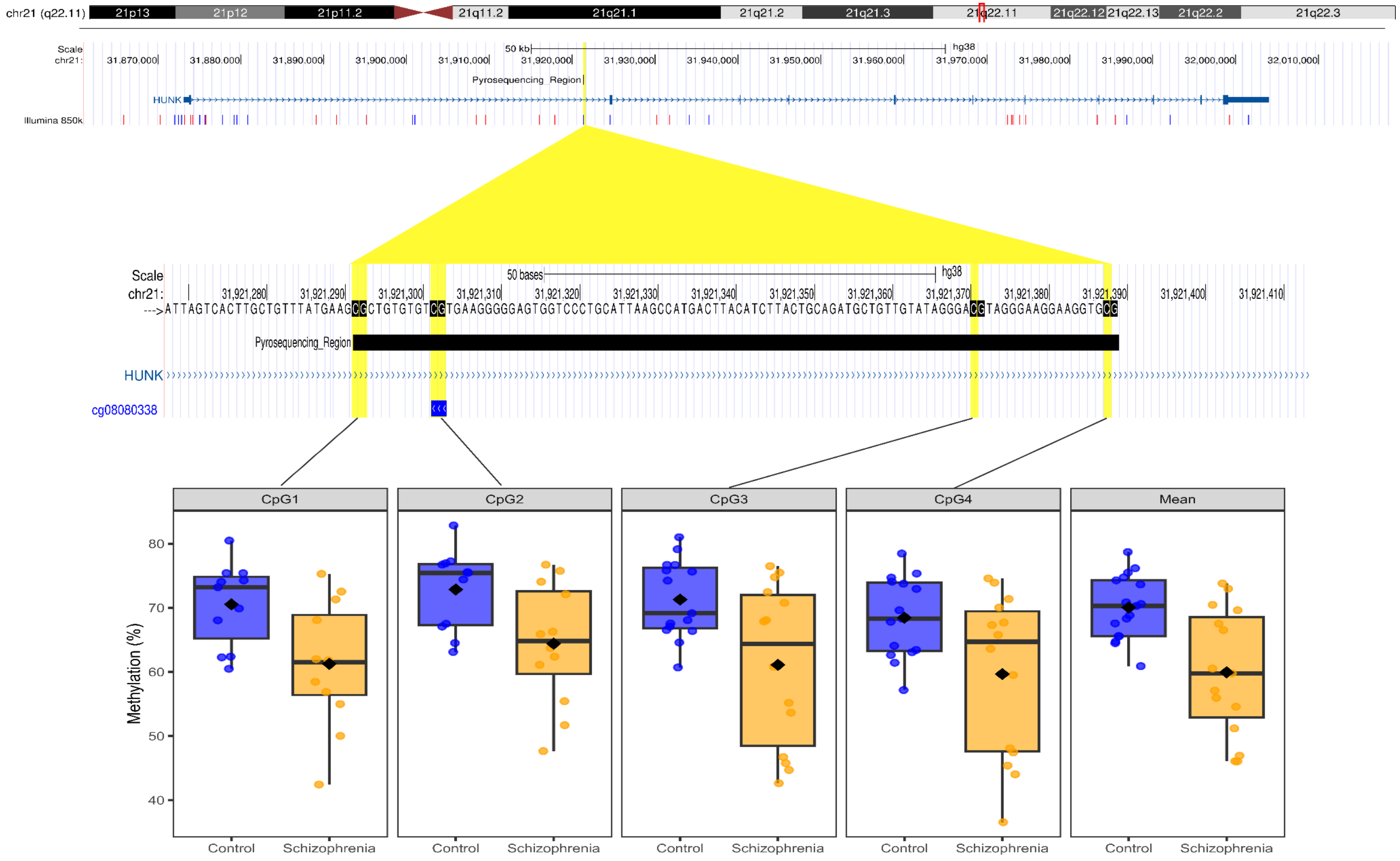

**Supplementary Figure 11: Genomic context of schizophrenia-associated DMPs located between *SSC5D* and *SBK2*.** Mini-Manhattan plot showing the location of two experiment-wide significant schizophrenia-associated DMPs, cg12421899 and cg25778661, within an intergenic region between *SSC5D* and *SBK2*. The two CpG sites are separated by only 14 bp and exhibit concordant schizophrenia-associated hypomethylation in neuronal nuclei. Tracks show local association statistics, annotated genes, Illumina EPIC array probes and neuronal enhancer regions from ENCODE.

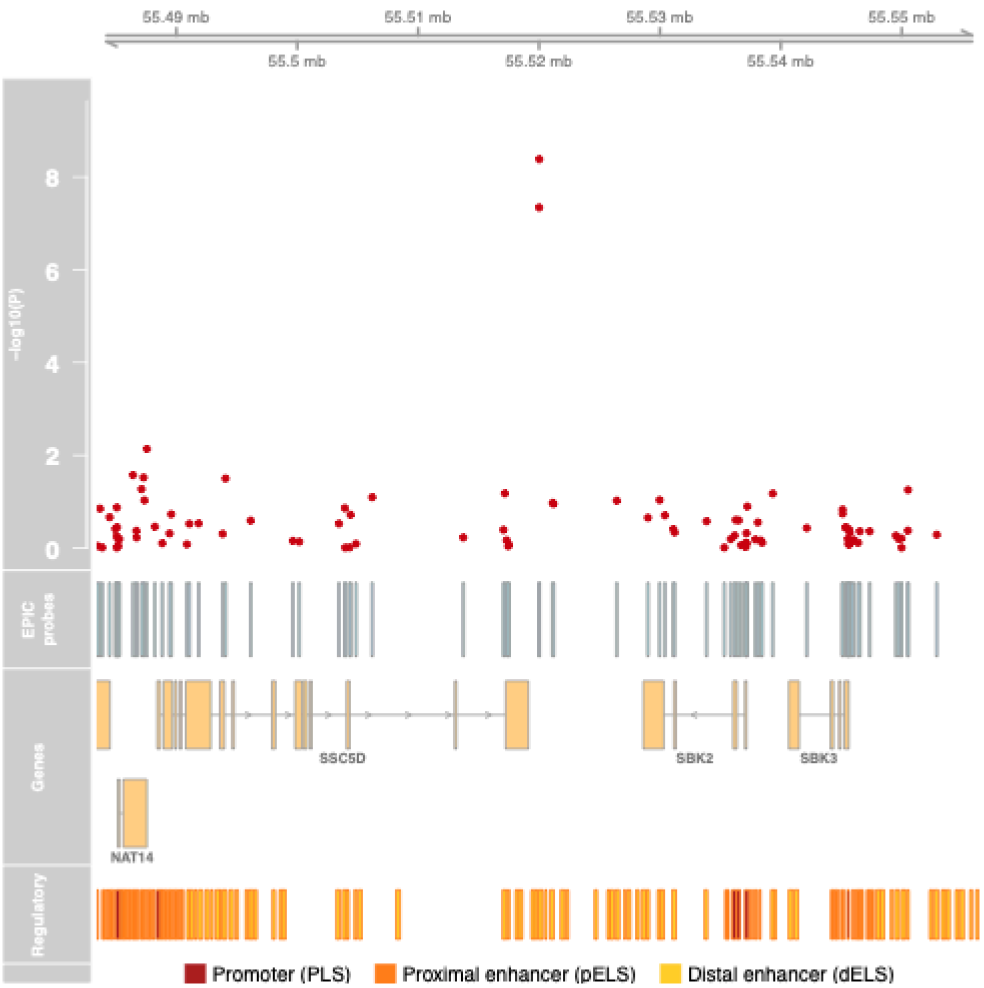

**Supplementary Figure 12: Schizophrenia-associated DNA methylation differences identified in NeuN+ nuclei from the prefrontal cortex show correlated effect sizes in NeuN+ nuclei isolated from additional brain regions.** Scatterplots comparing schizophrenia-associated DNA methylation effect sizes (case-control differences) for discovery-threshold neuronal DMPs identified in prefrontal cortex (PFC) NeuN+ nuclei with the corresponding effect sizes observed in neuronal nuclei isolated from a subset of matched donors (n = 20) (left panel), the hippocampus (middle panel) and striatum (right panel). The left panel compares effect sizes estimated in the full PFC neuronal cohort with those obtained from the matched PFC subsample, demonstrating the high representativeness of the selected subset. DMPs reaching experiment-wide significance in the full PFC neuronal EWAS are highlighted in red. Pearson's correlation coefficients (R) are shown for each comparison. These analyses demonstrate evidence for modest cross-region concordance of schizophrenia-associated neuronal DNA methylation differences, with stronger correspondence observed between PFC and hippocampal neurons than between PFC and striatal neurons.

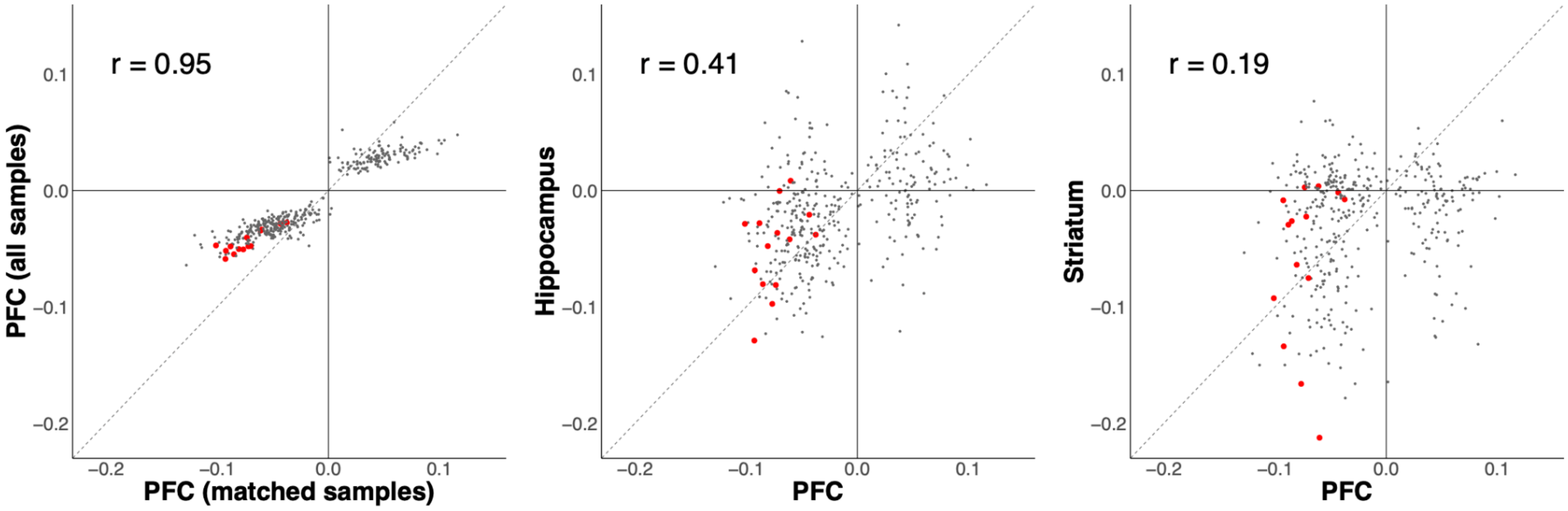

**Supplementary Figure 13: Regional heterogeneity of schizophrenia-associated neuronal DNA methylation differences across brain regions.** Boxplots showing DNA methylation levels at representative schizophrenia-associated neuronal DMPs annotated to HUNK (cg08080338, left) and SS5CD (cg12421899, right) in neuron-enriched (NeuN+) nuclei isolated from prefrontal cortex (PFC), hippocampus and striatum. Hypomethylation at the top-ranked HUNK-associated DMP was restricted to PFC neurons (matched PFC NeuN+ nuclei P = 0.000816, hippocampus NeuN+ nuclei P = 0.417, striatum NeuN+ nuclei P = 0.615). In contrast, the SS5CD-associated DMP showed consistent schizophrenia-associated DNA methylation differences in neuronal nuclei from all three brain regions (matched PFC NeuN+ nuclei P = 0.000278, hippocampus NeuN+ nuclei P = 0.0320, striatum NeuN+ nuclei P = 0.00413). Boxplots display median and interquartile range, with individual donor values overlaid.

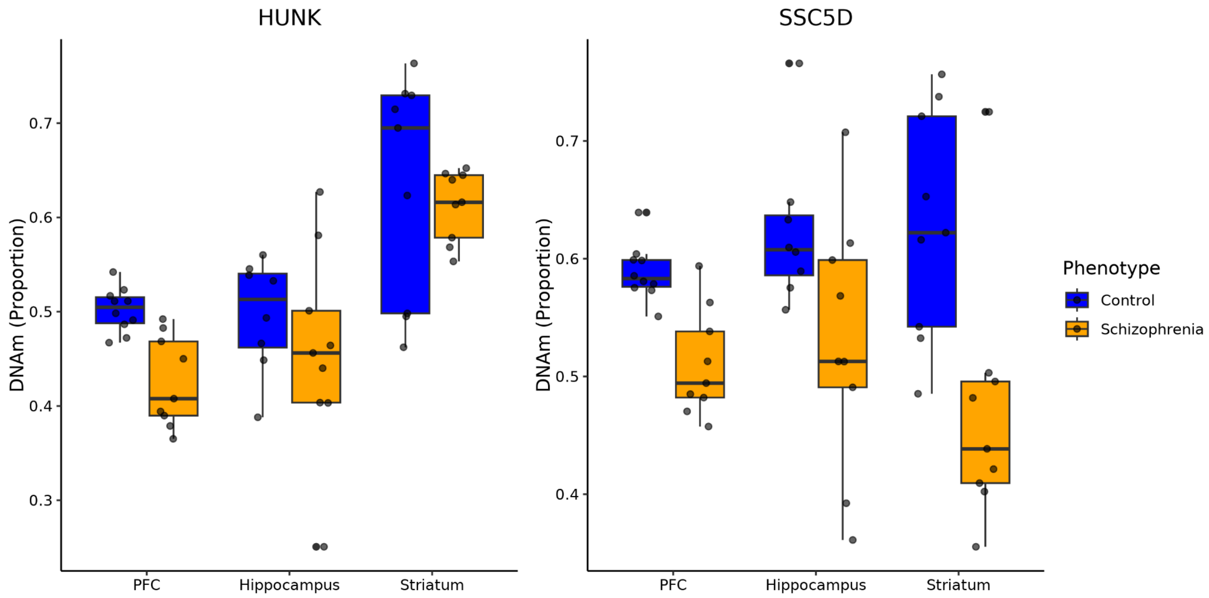
